# Why do some fungi want to be sterile? The role of dysfunctional Pro1 in the rice blast fungus

**DOI:** 10.1101/2023.01.23.525283

**Authors:** Momotaka Uchida, Takahiro Konishi, Ayaka Fujigasaki, Kohtetsu Kita, Tsutomu Arie, Tohru Teraoka, Takayuki Arazoe, Takashi Kamakura

## Abstract

Although sexual reproduction is widespread in eukaryotes, some fungal species can only reproduce asexually. Therefore, loss of sexual reproduction may confer survival advantages under certain conditions in certain species. In the rice blast fungus *Pyricularia* (*Magnaporthe*) *oryzae*, several isolates from the region of origin retain mating ability (female fertility), but most isolates are female sterile. Therefore, it is hypothesized that female fertility was lost during its spread from the origin to the rest of the world, and *P. oryzae* is an ideal biological model for studying the cause of the evolutionary shift in the reproductive mode. Here, we show that functional mutations of Pro1, a global transcriptional regulator of mating-related genes in filamentous fungi, is one cause of loss of female fertility in this fungus. Employing backcrossing between female-fertile and female-sterile field isolates, we identified the putative genomic region involved in female sterility by comparative genomics between the genomes of F_4_ female-fertile and -sterile progenies. Further genotyping, linkage, and functional analyses revealed that the functional mutation of Pro1 causes the loss of female fertility. RNA sequencing analysis showed that Pro1 regulates global gene expression, including that of several mating-related genes. The dysfunctional Pro1 did not affect the infection processes, such as conidial germination, appressorium formation, and penetration, but conidial release from conidiophores was increased. Furthermore, various types of mutations in Pro1 were detected in geographically distant *P. oryzae*, including pandemic isolates of wheat blast fungus. These results provide the first evidence that loss of female fertility may be advantageous to the life cycle of some plant pathogenic fungi.

**Significance:** Many pathogenic and industrial fungi are thought to have abdicated sexual reproduction, but the mechanisms and biological importance have been a long-standing mystery. Discovering why such fungi lost fertility is important to understand their survival strategies. Here, we revealed the genetic basis of how the rice blast fungus lost female fertility in nature and how this affects the life cycle. This has important implications for understanding evolution of blast pathogens and for developing an effective management strategy to control blast disease before a pandemic. Our findings also provide an additional perspective on advantages of asexual reproduction in some eukaryotes.

## Main Text

Sexual reproduction is common in eukaryotic organisms and is considered to have evolved once in the last eukaryotic common ancestor (*1, 2, 3*). The process of meiosis in mating can produce genetic diversity and purge deleterious mutations in the genome of the progeny (*4, 5*). However, this mode of reproduction might be associated with various costs and risks (*6, 7*); for example, finding mating partners is both time and energy expensive, and genetic shuffling with other individuals increases the risks of sexually transmitted diseases. Theoretical analyses suggest that the advantage of sexual reproduction is often quite restrictive, and switching to asexual reproduction within a sexual species may enable asexual progeny to outcompete and displace sexual progeny (*8, 9*). Thus, why sex is evolutionarily advantageous remains largely unexplored (paradox of sex).

Fungi and animals evolved from a common single-celled ancestor and constitute the opisthokonts (*10, 11*). Each fungal group shows a wide variety of sexual development, and most known fungal species reproduce both asexually and sexually (*10, 12*). Although fungal sexual reproduction may be cryptic or facultative, several industrial and pathogenic fungi are considered to reproduce only asexually in their life cycles owing to partial or complete sterility (*12–17*).

Interestingly, these asexual fungi maintain certain key components of the mating and meiosis genes, suggesting that evolutionary shifts from sexual to asexual reproduction have occurred and that asexual fungi have dominated by natural selection.

*Pyricularia oryzae* (syn. *Magnaporthe oryzae*) is a haploid filamentous ascomycete fungus and causes blast disease on a variety of cereal and grass hosts, such as rice (*Oryza sativa*), wheat (*Triticum aestivum*), barley (*Hordeum vulgare*), finger millet (*Eleusine coracana*), foxtail millet (*Setaria italica* and *S. viridis*), perennial ryegrass (*Lolium perenne*), annual ryegrass (*Lolium multiflorum*), southern cutgrass (*Leersia hexandra*), and goosegrass (*Eleusine indica*) (*18*). The rice blast fungus, which causes the most devastating disease of cultivated rice worldwide, is well studied as a model plant pathogenic fungus. The pathogen is indicated to have diverged from a *S. italica*-infecting strain in the Middle Yangtze Valley of China approximately 2500–7500 years ago, during or shortly after rice domestication in East Asia (*19*). The infection cycle of this fungus consists of asexual reproduction. The infection starts when a three-celled asexual spore (conidium) adheres to the hydrophobic surface of a rice leaf. The germinated conidium develops a dome-shaped infection-specific structure, termed an appressorium, to invade an epidermal cell (*20, 21*). After invasion, filamentous invasive hyphae spread to adjacent cells and develop a visible disease lesion on the leaf surface. In humid conditions, conidiophores and conidia are produced on the lesion, and the conidia are delivered to new host plants by wind or dewdrop splashes, initiating a new cycle (*20, 21*). In addition to the asexual life cycle, the sexual reproduction of *P. oryzae* has been observed under laboratory conditions (*22, 23*). *P. oryzae* is a heterothallic fungus and carries either *MAT1-1* (*MAT1-α*) or *MAT1-2* (*MAT1-HMG*) idiomorphs on chromosome 7 (*24, 25*). Sexual reproduction occurs by the interaction between two strains with the opposite mating type, and produces a multicellular female organ, termed a perithecium, in which asci develop (*22–26*) (Fig. 1A). Eight four-celled ascospores (sexual spores) are produced per ascus through meiosis and mitosis (*22–26*) (Fig. 1A). Sexual reproduction requires at least one female-fertile strain that is capable of producing perithecia; however, female fertility has been lost in the most field isolates of the rice blast fungus (*27–29*). Interestingly, sexual reproduction has been lost in some ascomycete plant pathogens, but the contributing factors and biological importance have long remained unknown (*30-32*).

**Fig. 1.**
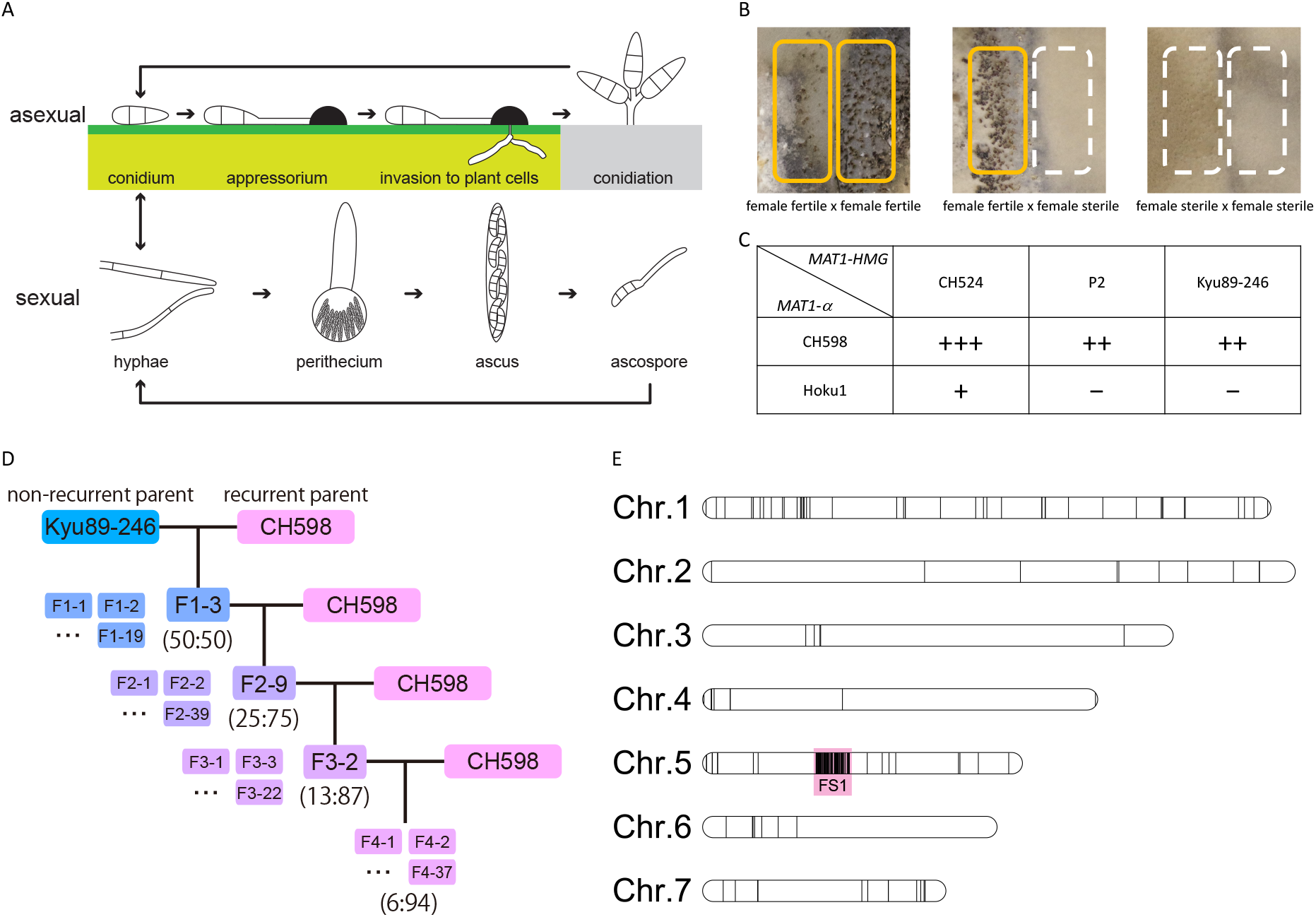
Perithecium formation, assessment of inheritance of female sterility, and nucleotide substitutions unique to female-sterile isolates in *Pyricularia oryzae*. (**A**) The life cycle of *P. oryzae*. (**B**) Perithecium formation in *P. oryzae*. Perithecia develop on the border of two strains, in two lines when both strains are female fertile (left) or in one line when one strain is female fertile and the other is female sterile (middle). No perithecia develop between two female-sterile strains (right). (**C**) Perithecia development in crosses between field isolates harboring *MAT1-HMG*. +++, >100 perithecia per plate; ++, >10 perithecia per plate; +, >0 perithecia per plate; −, no perithecia. (**D**) Schematic diagram of the recurrent backcrossing strategy between the field isolates Kyu89-246 and CH598. One female-sterile progeny of each generation harboring *MAT1-HMG* was crossed with CH598. Parentheses represent the theoretical percentage of inherited genomic content (Kyu89-246:CH598). (**E**) Nucleotide substitutions between female-fertile and -sterile progenies. Black bars in each chromosome represent loci with unique nucleotide substitutions in the genome of female-sterile progenies. The substitutions are concentrated in a region on chromosome 5, termed the Female Sterility 1 (FS1) region (red square).

Some isolates in the region of origin of *P. oryzae* retain female fertility (*27–29*), and population genetic/genomic studies have provided evidence that sexual recombination events continue to occur, at least in limited areas within this region (*27–29,33*). Therefore, it has been hypothesized that the loss of sexual reproduction occurred during its geographic spread from the region of origin. *P. oryzae* is an ideal biological model for studying the reason for the evolutionary shift in reproductive mode (*12, 27*). Although many mating-related genes have been identified in ascomycete fungi (*34–37*), the genes and mechanisms responsible for the loss of sexual reproduction in nature have remained unclear. In this study, we established that functional mutation of the transcriptional regulator Pro1 causes loss of female fertility and leads to increase in the release of conidia from conidiophores. In addition, various types of mutations in *Pro1* were independently detected in several geographically distant *P. oryzae* isolates. Given that the increase in conidial detachment would be effective for geographic spread and dispersal of conidia to other host plants, we provide experimental evidence that loss of sexual reproduction may confer a fitness advantage for the life cycles in plant pathogenic fungi.

## Results

### Generation of F_4_ near-isogenic strains by recurrent backcrossing

We set out to obtain female-sterile and female-fertile near-isogenic strains of *P. oryzae* employing a backcrossing strategy. First, we evaluated mating capability of rice-infecting field isolates: CH598 (*MAT1-α*) and CH524 (*MAT1-HMG*) collected from Yunnan, China (the region of origin of this pathogen), and Hoku-1 (*MAT1-α*; synonymous with Hoku1), P2 (*MAT1-HMG*; synonymous with P-2), and Kyu89-246 (*MAT1-HMG*) collected from Japan (25). When two strains are female-fertile and male-fertile, two lines of perithecia are formed, one for each female-fertile strain (Fig. 1B, C). Crosses of a female-fertile strain with a female-sterile strain develop one line of perithecia, whereas no perithecia are formed in crosses between two female-sterile strains (Fig. 1B, C). In our experiment, perithecia were developed in two lines by crossing CH598×CH524, whereas few and dotted perithecia were developed by crossing Hoku-1×CH524 (fig. S1A). One line of perithecia were formed by crossing CH598×Kyu89-246 and CH598×P2 (fig. S1A). No perithecia were formed by crossing Hoku-1×P2 and Hoku-1×Kyu89-246 (fig. S1A). Thus, CH598 and CH524 were defined as female-fertile strains, and Hoku-1, P2 and Kyu89-246 were defined as female-sterile strains (Fig. 1C). Mature asci containing viable ascospores were obtained by crossing CH598×CH524 and CH598×Kyu89-246, although viable ascospores were not frequent for CH598×Kyu89-246 (fig. S1B). Perithecia produced by Hoku-1×CH524 and CH598×P2 contained no ascospores (fig. S1B). From these results, CH598 and Kyu89-246 were subsequently used for the backcrossing analysis.

We obtained 91 F_1_ progenies (57 *MAT1-α* and 34 *MAT1-HMG*) from CH598×Kyu89-246. All 57 progenies harboring *MAT1-α* were backcrossed with Kyu89-246, and the six progenies produced perithecia (female fertility). However, the perithecia did not contain mature ascospores (fig. S2). In addition, by backcrossing with CH598, two out of 19 progenies harboring *MAT1-HMG* formed two lines of perithecia with mature ascospores (female fertility), and the other progenies formed one line (female sterility) (fig. S2). Thus, one female-sterile progeny harboring *MAT1-HMG* (F_1_-3) was used for subsequent backcrossing with CH598 (Fig. 1D). Each female-sterile F_2_ (F_2_-9) and F_3_ (F_3_-2) progeny harboring *MAT1-HMG* was crossed with CH598, and in total 37 F_4_ progenies were obtained (Fig. 1D and fig. S2). The segregation ratio of female-fertile and -sterile F_4_ progenies was approximately 1:1, independent of the *MAT1* locus (table S1), suggesting that the tested F_3_ progeny (F_3_-2) carried a single gene or genomic region involved in female sterility.

### Identification of genomic region responsible for loss of female fertility

To determine the gene or genomic region responsible for loss of female fertility, we conducted a comparative genomic analysis between the female-fertile and -sterile F_4_ progenies. Four independent genomic DNA extracts from each four female-fertile and four female-sterile F_4_ progenies were separately pooled and sequenced using an Illumina HiSeq platform. Theoretically, each F_4_ progeny inherits 6.25% (1/16) of the genome derived from Kyu89-246. In addition, by mixing four genomic DNAs, an eight-fold higher resolution is expected because each additional genome of each progeny doubles the resolution. Each read obtained from female-fertile and -sterile progenies was mapped to the *P. oryzae* reference genome, and 40,080 and 40,422 variants were detected from each set of reads. By comparison, 9,066 variants were detected in the genomes of female-sterile progenies. Visualizing of the loci of variants in the reference genome showed a variant-rich region (112 substitutions within 516 kb) on chromosome 5 (Fig. 1E). This region was designated as FS1 (Female Sterility 1) and was analyzed in greater detail to identify candidate genes involved in female fertility.

Among 168 protein-coding genes located in the FS1 region, amino acid substitutions were detected in 11 genes (fig. S3). Thus, we disrupted these candidate genes in CH598 using the CRISPR/Cas9 system (*38*) and evaluated female fertility. The deletion mutants for MGG_11498 (hypothetical protein) and MGG_00779 (choline dehydrogenase) showed decrease perithecia formation when crossed with Kyu89-246 (fig. S3). The deletion mutants for MGG_00722, MGG_00706, and MGG_00693, encoding hypothetical proteins, could not be obtained. Deletion of the remaining six genes, MGG_00791, MGG_00771, MGG_00747, MGG_17384, MGG_11512, and MGG_17398, resulted in no remarkable changes in mating ability (fig. S3). To test whether the five genes MGG_11498, MGG_00779, MGG_00722, MGG_00706, MGG_00693 were responsible for loss of female fertility in Kyu89-246, we introduced female-sterile-type amino acid mutations into CH598 by CRISPR/Cas9-mediated base editing (*39*).

None of these mutations affected female fertility, indicating that these genes are not involved in female sterility.

Because genes responsible for loss of female fertility were not located within the FS1 region, we conducted linkage analysis to clarify whether the FS1 and neighboring regions are associated with female sterility. A *hygromycin phosphotransferase* (*HPH*) gene cassette was knocked-in to each left (L), central (C), and right (R) locus of the FS1 region in CH598 using the CRISPR/Cas9 system (Fig. 2A). Each obtained transformant (CH598 FS1L-*HPH*, CH598 FS1C-*HPH*, or CH598 FS1R-*HPH*) was crossed with the female-sterile F_4_-5 (*MAT1-HMG*) and more than 50 ascospores were randomly collected from each cross. Most of the *HPH*-possessing progenies were female fertile, and the FS1R locus showed slightly stronger linkage than the other loci (Fig. 2A). However, several female-sterile *HPH*-possessing progenies were obtained in all crosses, suggesting that the responsible gene was located outside of this region (Fig. 2A).

**Fig. 2.**
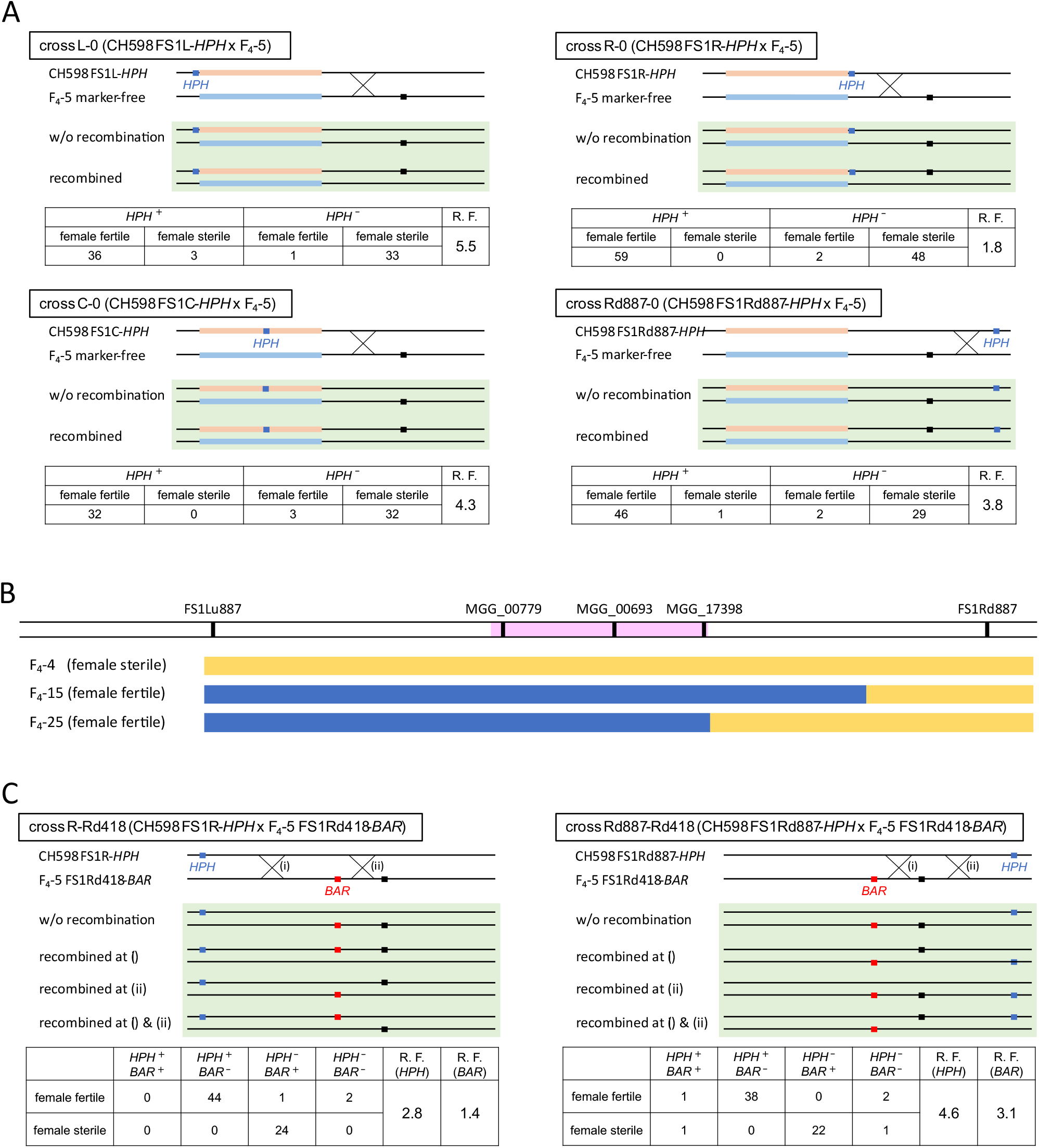
Linkage and genotyping analyses. (**A**) Linkage analysis between the integrated *hygromycin phosphotransferase* (*HPH*) genes and female sterility. The marker-integrated CH598 transformants were crossed with marker-free F_4_-5. In each cross, genetic maps of the parental genomes are presented at the top, and those of possible recombinant genomes of progenies are presented in the green box. Light red and light blue bars indicate each FS1 region of CH598 and Kyu89-246 (F4-5), respectively. Black and blue boxes represent the loci of putative female sterility gene and *HPH* gene, respectively. *HPH*^+^, hygromycin resistance; *HPH*^-^, hygromycin sensitive; R.F., recombination frequency (%) between *HPH* and the female sterility. (**B**) Determined genotypes and phenotypes of F_4_ progenies. Schematic representation of the loci of each target sequence for genotyping is presented at the top. Bars shaded with pink, yellow, and blue represent the FS1 region, female-fertile genotype, and female-sterile genotype, respectively. (**C**) Linkage analysis between integrated *BAR* genes and the female sterility. Light red and light blue bars indicate each FS1 region from CH598 and Kyu89-246, respectively. Black, blue, and red boxes represent the loci of putative female sterility gene, *HPH* gene, and *BAR* gene, respectively. *HPH*^+^, hygromycin resistance; *HPH*^-^, hygromycin sensitive; *BAR*^+^, bialaphos resistance; *BAR*^-^, bialaphos sensitive; R.F. (*HPH*), recombination frequency (%) between *HPH* and the female sterility; R.F. (*BAR*), recombination frequency (%) between *BAR* and the female sterility.

In addition to the linkage analysis, we analyzed genotypes around the FS1 region in the F_4_ progenies to examine whether the DNA sequences were consistent with the phenotype. SNPs within MGG_00779, MGG_00693, and MGG_00773 genes were selected as targets for genotyping because these genes were close to the FS1L, FS1C, and FS1R loci, respectively (Fig. 2A and B). Among 37 F_4_ progenies, 34 had the SNPs corresponding to the phenotype, but F_4_-15 (female fertile), F_4_-25 (female fertile), and F_4_-4 (female sterile) lacked the SNPs (Fig. 2B). These results were consistent with the linkage results, which contained *HPH*^+^ female-sterile and *HPH*^-^ female-fertile progenies (Fig. 2A). Thus, we analyzed SNPs located outside of the FS1 region in F_4_-15, -25, and -4. The SNPs located 887 kb upstream (within the putative promoter region of MGG_17324) of FS1L (FS1Lu887) were inconsistent with their phenotypes (Fig. 2B). In contrast, the SNPs located 887 kb downstream from FS1R (within MGG_00399 ORF) (FS1Rd887) were correlated to the phenotype (female fertility) in F_4_-15 and -25 (Fig. 2B). These results strongly suggested that the responsible gene was located downstream of the FS1 region. However, F_4_-4 (female sterility) contained the female-fertile-type SNPs at the FS1Rd887 locus (Fig. 2B). Therefore, we speculated that F_4_-4 was a spontaneous mutant that lost female fertility during laboratory cultivation, similar to previously reported female-sterile-evolved strains (*12*; see Discussion and fig.S4).

To determine the genetic region involved in loss of female fertility, the *HPH* cassette was knocked-in at FS1Rd887 in CH598 (Fig. 2A). Crossed with the female-sterile F_4_-5 (*MAT1-HMG*), the transformant (CH598 FS1Rd887-*HPH*) showed a recombination frequency of 3.8% (Fig. 2A). This frequency was higher than that in FS1R (1.8%) and lower than that in FS1L (5.5%) and FS1C (4.3%) (Fig. 2A), indicating that the region involved in female sterility was located between FS1R and FS1Rd887, and was closer to FS1R in genetic distance. We inserted the *bialaphos resistance* (*BAR*) gene cassette at 418 kb downstream from FS1R (FS1Rd418) in the female-sterile F_4_-5 progeny (Fig. 2), and the transformant (F_4_-5 FS1Rd418-*BAR*) was crossed with CH598 FS1R-*HPH* or FS1Rd887-*HPH*. The recombination frequencies of *BAR* were lower than those of *HPH*, indicating that the female sterility gene was more strongly linked to FS1Rd418 than FS1R or FS1Rd887 (Fig. 2C). Furthermore, the two progenies in which recombination occurred between FS1R and FS1Rd418 (*HPH*^*−*^ */BAR*^*−*^ or *HPH*^*+*^*/BAR*^*+*^) were female fertile, whereas five progenies with recombination between FS1Rd418 and FS1Rd887 (*HPH*^*−*^ */BAR*^*−*^ or *HPH*^*+*^*/BAR*^*+*^) contained both female-fertile and female-sterile strains (Fig. 2C), indicating that the gene responsible for loss of female fertility was located between FS1Rd418 and FS1Rd887. In addition, a consistent genotype was detected across this region in F_4_-15 and F_4_-25. From these results, we re-aligned the HiSeq reads of the female-fertile F_4_ progenies to the *de novo* assembled genome of female-sterile F4 progenies and re-mapped the detected substitutions to the reference genome. The cluster of nucleotide substitutions was isolated until 877 kb downstream of FS1 (fig. S5).

### Functional mutation of putative transcriptional regulator Pro1 induced loss of female fertility in Pyricularia oryzae Kyu89-246

Three genes (MGG_00494, MGG_14683, and MGG_00428) with amino acid substitution(s) in the female-sterile progenies were located within the FS1Rd418–FS1Rd887 region in the 70-15 genome (Table 1). Phenotypic analysis of deletion mutants of these genes in CH598 revealed that MGG_00494, which codes a putative Zn(II)_2_Cys_6_ zinc cluster transcriptional factor Pro-1 (hereafter *Pro1*), is necessary for perithecium formation (Fig. 3A). Complementation with the *Pro1* sequence derived from CH598 (female fertile) restored the fertility of the *∆pro1* mutant (Fig. 3A). The amino acid sequence similarity of Pro1 derived from CH598 (Pro1^CH598^) with *Sordaria macrospora* Pro1 (SMAC_00338) and *Neurospora crassa* Adv-1 (NCU07392) was 71.19% and 72.19%, respectively (fig. S5). Compared with Pro1^CH598^, two mutations, S16W and frameshift after G125 (8 bp deletion), which leads to protein truncation (Fig. 3B), were detected in the *Pro1* sequence of Kyu89-246 (*Pro1*^Kyu89-246^). To clarify whether these mutations affect Pro1 function, we generated three types of mutants in CH598: S16W (PM^S16W^), truncation (PM^truncation^), and S16W + truncation (PM^Kyu89-246^) (Fig. 3B). In crosses with Kyu89-246, PM^S16W^ and the wild type formed perithecia, whereas PM^truncation^ and PM^Kyu89-246^ did not form perithecia (Fig. 3A). These results showed that the truncation type mutation (loss of function of Pro1) induces loss of female fertility in CH598.

**Table 1.** Candidate genes between the FS1Ed418 and FS1Ed887 regions.

| Locus | Protein | Distance from FS1D<br>(kb downstream) | Amino acid substitutions |
| --- | --- | --- | --- |
| MGG_00494 | transcriptional regulatory protein Pro-1 | 719 | S16W and frameshift after G125 |
| MGG_14683 | hypothetical protein | 786 | P494A |
| MGG_00428 | conidial yellow pigment biosynthesis polyketide synthase | 806 | V681L and T866A |

**Fig. 3.**
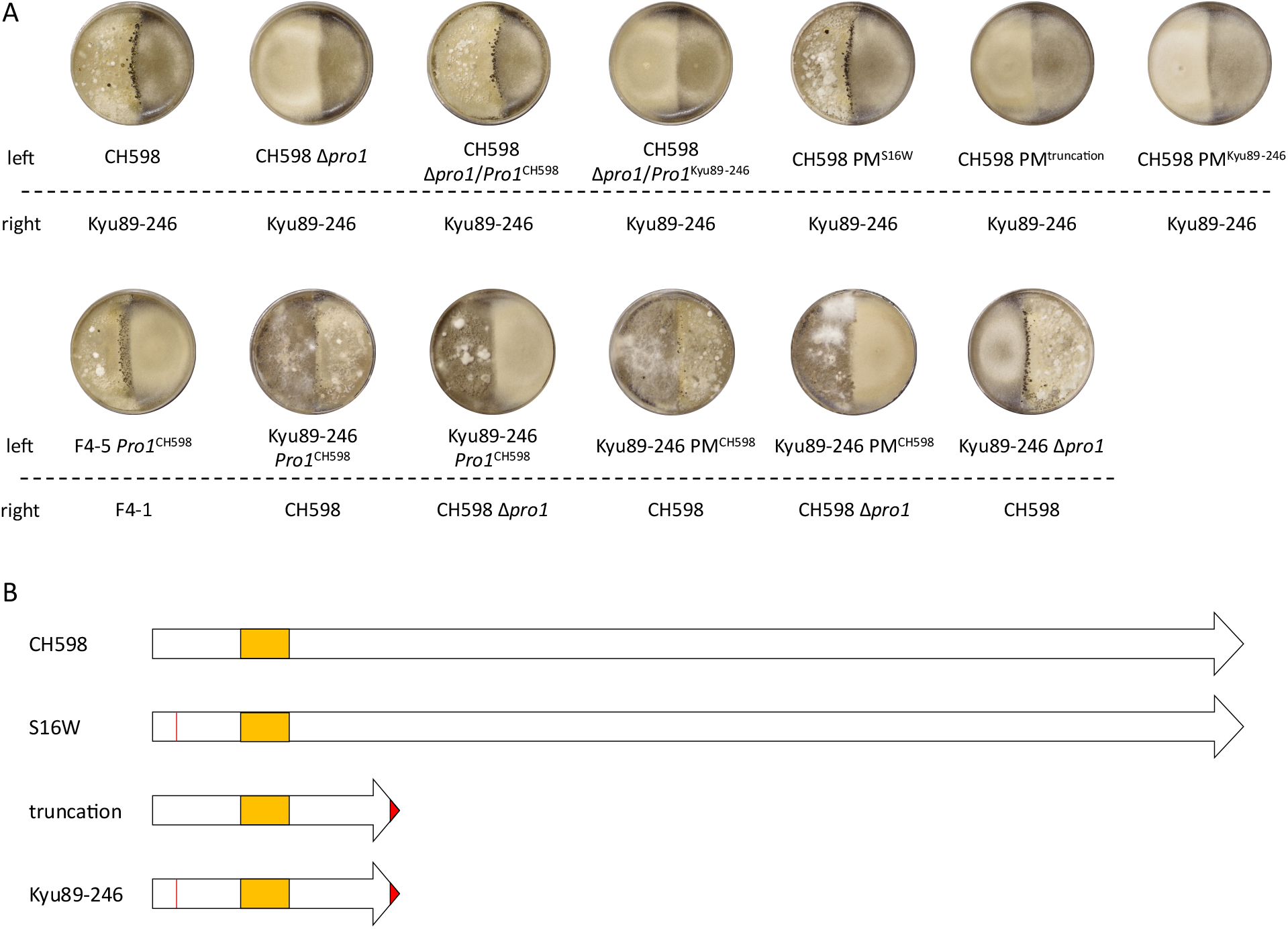
Mating capabilities of *Pro1*-defiicient mutants. (**A**) Female fertility for CH598, F_4_-5, Kyu89-246, and their transformants. (**B**) Schematic diagram of Pro1 mutations in the parental isolates (CH598 and Kyu89-246) and transformants (PM^S16W^ and PM^trancation^). Regions colored red represent the substituted amino acid sequence. PM^S16W^ possesses the S16W mutation. PM^truncation^ possesses the truncation type mutation (frameshift after G125). PM^Kyu89-246^ has both S16W and truncation type mutations. Orange boxes indicate Zn(II)_2_Cys_6_ DNA-binding domain.

To validate whether the functional Pro1 restores the female fertility in female-sterile field isolates, we introduced *Pro1* derived from CH598 (*Pro1*^CH598^) into F_4_-5 and Kyu89-246. As a result, the F_4_-5 transformant (F_4_-5 *Pro1*^CH598^) showed fully restored female fertility (Fig. 3A), indicating that *Pro1* is one gene responsible for loss of female fertility. This result is consistent with the segregation ratio (1:1) of female fertility and female sterility in the F_4_ progeny.

Meanwhile, recovery of female fertility in Kyu89-246 *Pro1*^CH598^ was not observed (Fig. 3A). Thus, we directly modified the mutated *Pro1* in Kyu89-246 (*Pro1*^Kyu89^) by base editing, to replace it with CH598-type *Pro1* (PM^CH598^) (Fig. 3B); however, the functional *Pro1* sequence in Kyu89-246 could not restore female fertility. These results suggested that more than one gene is responsible for female sterility in the isolate Kyu89-246.

### Pro1 functioned as a transcriptional regulator and regulated mating-related genes

Both *S. macrospora* Pro-1 and *N. crassa* Adv-1 are reported to play a role as a transcriptional regulator involved in sexual signal transduction and perithecium formation (*40– 42*). To validate the Pro1 function in *P. oryzae*, we performed RNA sequencing (RNA-seq) analysis using the two parental strains (CH598 and Kyu89-246), three female-fertile F_4_ progenies harboring *Pro1*^CH598^, and three female-sterile F_4_ progenies carrying *Pro1*^Kyu89-246^. Principal component analysis revealed that the mycelial transcriptome on RY media (supernatant of rice flour broth) was clustered into three groups: 1. Female-fertile strains including CH598, 2.

Female-sterile progenies, and 3. Kyu89-246 (Fig. 4A). The group of female-sterile progenies was closer to female-fertile strains than Kyu89-246; therefore, this grouping may reflect the genetic background of each strain. In addition, a two-dimensional heatmap revealed similar clustering (Fig. 4B).

**Fig. 4.**
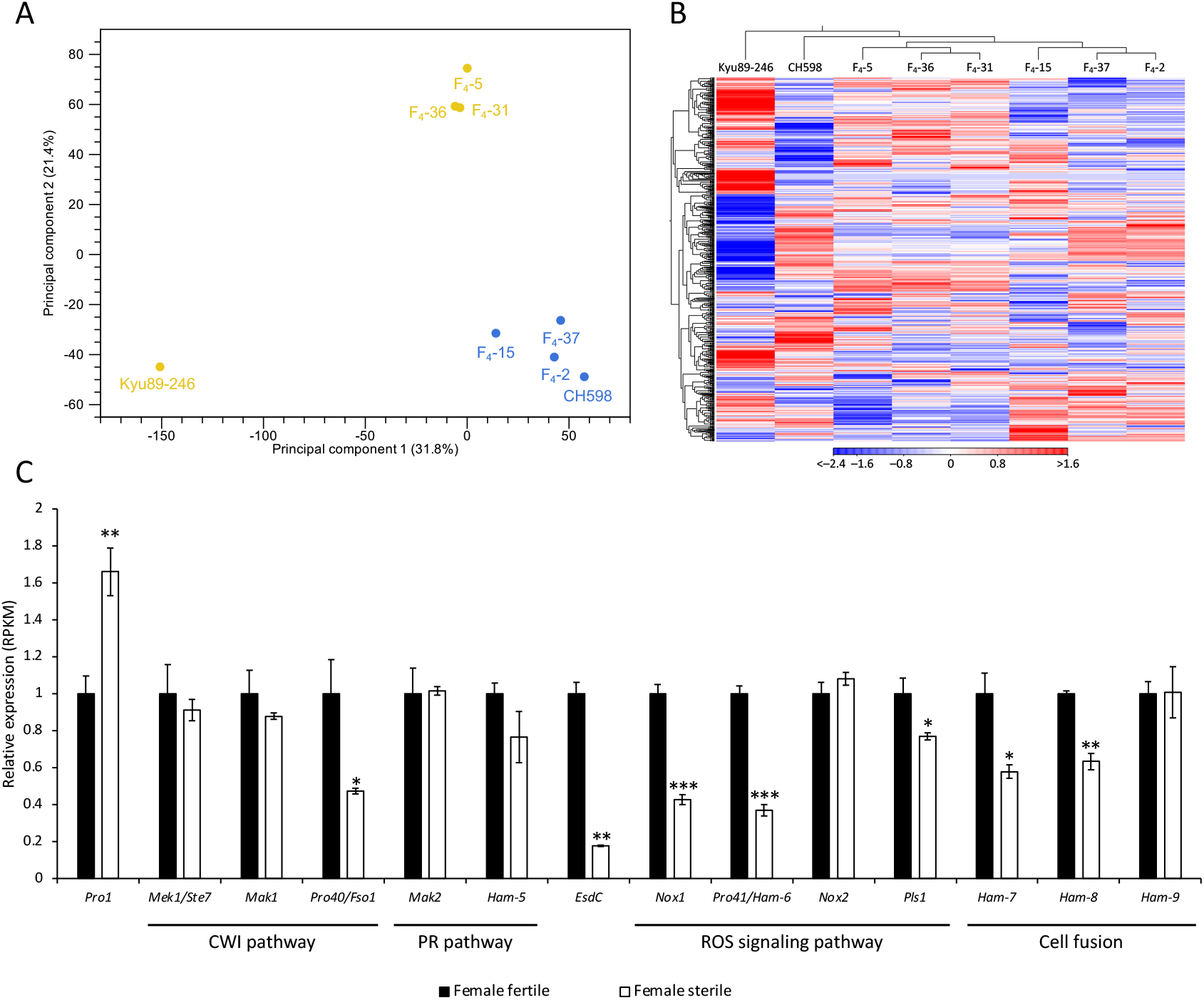
Transcriptomic analysis of F_4_ progenies. (**A**) Principal component analysis of RNA-seq based expression. The expression profiles of female-fertile progenies and CH598 (possessing functional Pro1) are colored with blue and of female-sterile progenies and Kyu-89-246 (with non-functional Pro1) are colored with yellow. (**B**) Expression patterns are shown in two-dimensional heatmap with Euclidean distance. Higher expression values are displayed red, and lower, blue. The female-fertile and female-sterile progenies were grouped separately. (**C**) Expression levels of some genes involved in cell wall integrity (CWI) pathway, pheromone response (PR) pathway, reactive oxygen species (ROS) signaling pathway, and cell fusion. Black and white bars indicate the expression levels of female-fertile and female-sterile progenies, respectively. Expression levels of each gene were calculated by dividing RPKM values in each strain by the average value of female-fertile strains. Bars and error bars indicate mean ± SEM. \**p* < 0.05, \*\**p* < 0.01, \*\*\**p* < 0.001 (Welch’s *t*-test).

As well as the *∆pro1* mutant of *S. macrospora* (*42*), the RNA-seq analysis showed that the expression of genes involved in the early stage of sexual development in *Aspergillus nidulans* (*EsdC*) (*43*), cell wall integrity pathway (*Pro40*) (*44*), hyphal fusion (*Ham-7* and *Ham-8*) (*45*), and reactive oxygen species signaling (*Nox1* and *Pro41*) (*46, 47*) were decreased in the female-sterile progenies (Fig. 4C). In addition to these genes, we identified 48 upregulated and 53 downregulated genes in the female-fertile progenies (fold change > 2 and *p* < 0.01; table S2).

Among the upregulated genes, 37 were annotated as hypothetical protein, and 17 were unique to *P. oryzae*. Annotated functions of other genes were highly diverse and no significant GO enrichment was detected (g:SCS algorithm in g:Profiler (*48*), *p* < 0.05). The 40 downregulated genes encoded hypothetical proteins, a half of which was unique to *P. oryzae* (table S2). These results indicated that Pro1 regulates several mating-related genes; therefore, loss-of-function of Pro1 leads to loss of female fertility in *P. oryzae*.

### Loss-of-function of Pro1 increases conidial release in P. oryzae

Given that most rice-infecting field isolates of *P. oryzae* show female sterility (*27–29*), we hypothesized that loss of female fertility caused by functional mutation of Pro1 provides fitness advantages in the asexual infection and saprotrophic life cycles. To test this hypothesis, we assessed the vegetative growth, conidiation, appressorium formation, and penetration in the *Pro1* mutants. Compared with CH598, the *∆pro1* mutant showed a slightly decreased growth rate on the rice flour medium (Fig. 5A), whereas the rate was comparable to CH598 on the complete medium (CM; fig. S7). Conidia production by the mutant was increased on rice flour medium (Fig. 5B) but was reduced on CM (fig. S7). Consistently, in CH598 ∆*pro1*/*Pro1*^Kyu89-246^, PM^truncation^, and PM^Kyu89-246^, the phenotypes were similar to that of the *∆pro1* mutant (Fig. 5A, B). We also examined the Pro1 function in Kyu89-246. PM^CH598^ grew more rapidly than Kyu89-246 on rice flour medium (Fig. 5A) but the growth rate was comparable to Kyu89-246 on CM (fig. S7). The number of conidia produced by PM^CH598^ was comparable to that of Kyu89-246 on rice flour medium (Fig. 5B) but was reduced on CM (fig. S7). Thus, Pro1 regulation of vegetative growth and conidiation in *P. oryzae* was dependent on nutrient conditions and/or strains. The rates of germination, appressorium formation, and penetration were not significantly different between the wild-type strains and mutants (fig. S8).

**Fig. 5.**
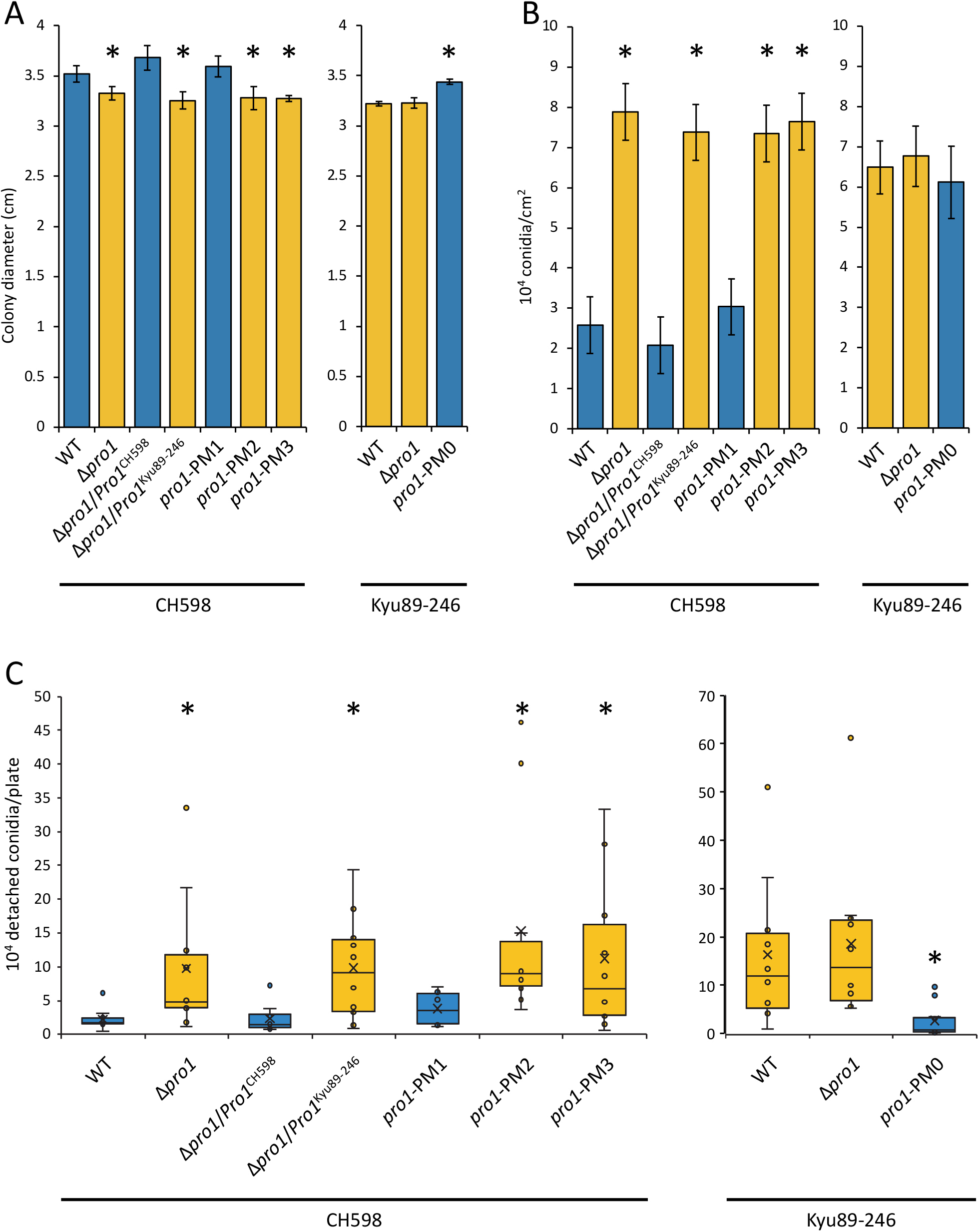
Asexual phenotypes affected by Pro1 function. (**A**) Colony diameter of 6-day-old cultures in the CH598 (left) and Kyu89-246 (right) genetic backgrounds on rice flour medium. Blue bars indicate strains possessing functional Pro1. Yellow bars indicate strains possessing dysfunctional Pro1. Diameters were measured in triplicate and repeated three times. Bars and error bars indicate mean ± SEM. (**B**) Number of conidia produced in the CH598 (left) and Kyu89-246 (right) genetic backgrounds on rice flour medium. Blue bars indicate strains possessing functional Pro1. Yellow bars indicate strains possessing dysfunctional Pro1. The experiment was repeated three times. Bars and error bars indicate mean ± SEM. (**C**) Box plots representing the number of conidia released per plate of rice flour medium in the CH598 (left) and Kyu89-246 (right) genetic backgrounds. The experiment was repeated ten times. Crosses indicate the mean, and the line within the box indicates the median. Blue boxes indicate strains possessing functional Pro1. Yellow boxes indicate strains possessing dysfunctional Pro1. * *p* < 0.05 (Welch’s *t*-test, compared with the corresponding wild type).

Because it has been previously reported that female-sterile-evolved strains increased conidial detachment (*12*), we further analyzed that in the *Pro1* mutants. Regarding CH598 and its mutants, detachment was approximately 4–7 times higher in the *∆pro1*, PM^truncation^, and PM^Kyu89-246^ mutants than in CH598, CH598 ∆*pro1*/*Pro1*^CH598^, and PM^S16W^ on rice flour medium (Fig. 5C). Similarly, for Kyu89-246, detachment in PM^CH598^ was six times fewer than that in the wild type and the *∆pro1* mutant (Fig. 5C). The increase in release of conidia in the *Pro1* mutants was observed on CM in both strains (fig. S7), suggesting that conidial release was not dependent on nutrient conditions and strain, which may confer an advantageous trait for propagation and/or colonization in the life cycle of *P. oryzae*.

### Mutations in Pro1 are widely distributed in field isolates of P. oryzae

To examine whether the mutations of *Pro1* are common in female-sterile isolates of *P. oryzae*, we first sequenced *Pro1* in the Japanese female-sterile field isolates P2 and Hoku-1. Both P2 and Hoku-1 had frameshift mutations after V605 (35 bp insertion: #2 mutation type in Fig. 6) and D317 (5 bp deletion and 43 bp insertion: #3 mutation type in Fig. 6) in the *Pro1* open reading frame. Similar to Kyu89-246, direct modification of the mutated *Pro1* did not restore female fertility in these isolates, whereas these mutations caused loss of female fertility in CH598 (fig. S8). These results indicated that P2 and Hoku-1 harbor other genes responsible for female sterility as well as Kyu89-246. We examined publicly available genome data for *P. oryzae*, including isolates from cereal and grass hosts. Various mutations in *Pro1* were detected in 137 of 329 genomic data sets. The mutations of *Pro1*, including those of Kyu89-246, P2, and Hoku-1, were classified into 29 variants, comprising amino acid substitutions, truncation, and mutation in the Zn(II)_2_Cys_6_ DNA-binding domain (Fig. 6 and table S3). Although many mutations were classified into variant #4, the same as the PM^S16W^ mutation which retains the Pro1 function, approximately two-thirds (20/29) of all variants showed truncation-type mutations. Given that C-terminal truncation of Pro1 detected in P2 (variant #2) caused female sterility in CH598 (fig. S9), the other truncation-type variants will also lead to loss-of-function of Pro1. The protein structural prediction of these variants by ColabFold (*49*) suggested that all of the variants alter the structure or folding of Pro1 (fig. S10). Therefore, the variants with amino acid substitutions (#5–12) may also cause loss-of-function of Pro1 or decrease in activity. The mutations, except for #4, were detected in geographically distant rice-infecting isolates collected from Italy, South Korea, China (Jiangsu), Suriname, and Japan, and some of them were phylogenetically distant despite close phylogenic relationships (fig. S11). These results suggest that loss of female fertility occurred independently during range expansion. In addition to the rice-infecting isolates, truncations of Pro1 were detected in the isolates from *T. aestivum, L. hexandra, S. viridis*, and *L. multiflorum*, suggesting that loss of female fertility caused by functional mutation of Pro1 is not uncommon in *P. oryzae* (table S3). Interestingly, the truncation-type mutations were frequently detected in pandemic isolates of the wheat blast fungus from Bangladesh (*50–52*)(#25 and #26 mutation types in Fig. 6 and table S3). Because the pathogen was first detected in Brazil and spread subsequently to the countries, loss of female fertility may be proceeding in the wheat blast fungus as well as the rice blast fungus.

**Fig. 6.**
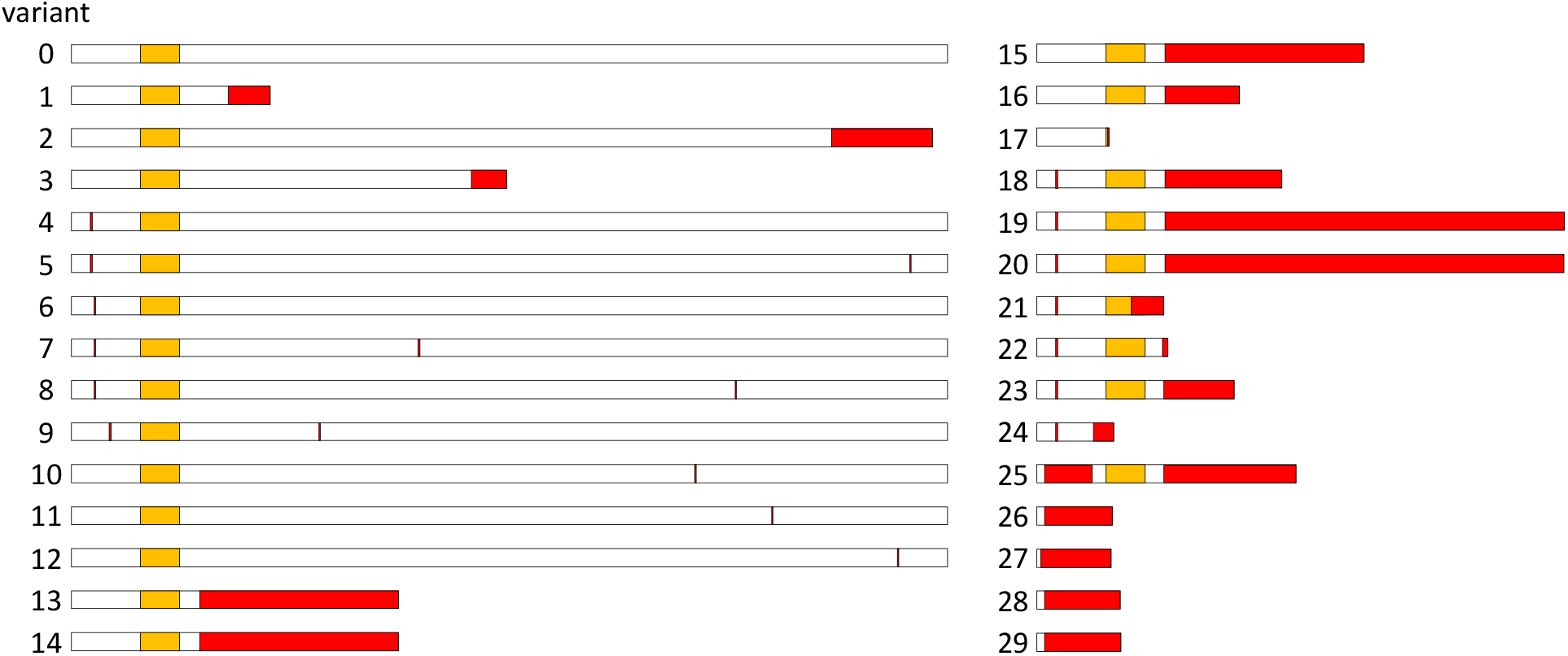
Pro1-variants detected in various assemblies deposited in NCBI and DDBJ. Mutations are shown in red. Orange boxes indicate Zn(II)_2_Cys_6_ DNA-binding domain. Pro1-variants #0, #1, #2, and #3 were detected in CH598, Kyu89-246, P2, and Hoku-1, respectively. The majority of Pro1-variants was variant #4, which possesses the S16W (PM^S16W^) mutation. Among the other Pro1-variants, 17 (#13-29) possessed truncation type mutations, and seven (#17, 21, 24, 26-29) possessed mutations which lead to loss of the Zn(II)_2_Cys_6_ DNA-binding domain.

## Discussion

Complementation of the mutated *Pro1* with the functional sequence in the female-sterile F_4_ progeny restored female fertility; however, fertility was not restored in the female-sterile field isolates Kyu89-246, P2, and Hoku-1 (fig. S9). These field isolates may contain at least one additional gene responsible for female sterility. The ratio of female-sterile and female-fertile F_1_ progenies was approximately 7:1 (female sterile: 68, female fertile: 8) and that of F_2_ progenies was approximately 3:1 (female sterile: 29, female fertile: 10). Consistently, the segregation ratio in F_1_ progenies possessing functional *Pro1*^CH598^ was approximately 3:1 (female sterile: 5, female fertile 2), and that in F_2_ progenies was approximately 1:1 (female sterile: 9, female fertile: 10). Although the number of progenies was insufficient to determine the segregation ratio, these results suggest that Kyu89-246 harbors three genes responsible for loss of female fertility, and the F_2_ progeny tested in this study carried two such genes.

The loss of Pro1 function in CH598 induced loss of female fertility (Fig. 3A). This phenotype was consistent with many sexual filamentous fungi, such as *S. macrospora* (*40*), *N. crassa* (*41*), *A. nidulans* (*53*), *Fusarium graminearum* (*54*), and *Cryphonectria parasitica* (*55*). As in *S. macrospora* (*42*), RNA-seq analysis showed that the expression levels of several mating-related genes (*Pro40, EsdC, Ham-7, Ham-8, Nox1, Pro41*, and *Pls1*) were decreased in the F_4_ progenies harboring *Pro1*^Kyu89^, compared with those in the progenies carrying *Pro1*^CH598^ (Fig. 4C). These reports and our findings provide that the role of Pro1 in perithecium formation is widely conserved in many sexual filamentous fungi, although the molecular mechanism of mating in *P. oryzae* remains poorly understood. In contrast to perithecium formation, the infection-related germination, appressorium formation, and penetration rates were not significantly affected by Pro1 function. These results were consistent with previous findings that the pathogenicity of a *Pro1* mutant is comparable to that of the wild-type strain (*56*). Because both female-fertile and -sterile parental isolates retain pathogenicity to rice, it is reasonable to consider that the infection process and pathogenicity are not affected by functional mutation of Pro1.

The conidial detachment was increased by loss of Pro1 function in CH598 and Kyu89-246 (Fig. 5C), which would provide survival advantages for their spread and propagation.However, to the best of our knowledge, molecular mechanisms of conidial release in filamentous fungi have not been elucidated. RNA-seq analysis showed that Pro1 regulates 77 uncharacterized genes encoding a hypothetical protein (table S2), and these may include novel functions associated with female fertility and conidial detachment. Female fertility in field isolates of *P. oryzae* from South China was rapidly lost following 10–19 rounds of subculture (*12*).Interestingly, these female-sterile strains, termed female-sterile-evolved strains, also show increased conidial release (*12*). Because the evolved strains were obtained by subculturing conidia, the increased ability for conidial detachment may be positively selected by the experimental procedure. Although the genes responsible in these strains have yet to be revealed, the efficient release of conidia and functional mutations of several genes involved in female fertility may have a trade-off relationship.

We also observed that loss of female fertility frequently occurred in CH598 under culture. Thus, we speculated that F_4_-4, in which the genotype of the FS1 region was inconsistent with the phenotype, is a female-sterile-evolved strain obtained under *in vitro* culture conditions. We confirmed whether loss of female fertility in the three female-sterile-evolved strains was caused by *Pro1* mutations, but no mutations were detected in these strains (data not shown). Principal component analysis showed that the transcription pattern of F_4_-4 was most similar to that of female-fertile progenies, rather than that of female-sterile progenies (fig. S4). Therefore, we concluded that the loss of female fertility in F_4_-4 was caused by the mutation of other genes.

From these findings, the loss of female fertility in *P. oryzae* may occur rapidly and frequently by mutations under various conditions and selective pressures.

Diverse types of *Pro1* mutation were independently detected in geographically distant isolates derived from various hosts (Fig. 6, fig. S11and table S3), consistent with the hypothesis that loss of female fertility occurred independently during or after the spread of this pathogen (*33*). Female sterility caused by Pro1 mutations may often occur, accompanied by pathogen dispersal. Interestingly, truncation-type mutations in *Pro1* have been frequently detected in pandemic isolates of the wheat blast fungus from Bangladesh (#25 and #26 in Fig. 6 and table S3). A clonal lineage (B71 lineage) of the wheat blast fungus has recently spread to Bangladesh following its independent introduction from South America (*52*). Importantly, the B71 isolate derived from Bolivia and clustered in the B71 lineage contained no *Pro1* mutations. This strongly suggests that loss of female fertility occurred after the introduction of this pathogen. In addition, genetic diversity among the isolates in the B71 lineage is reduced in comparison with South American isolates (*52*). This result is consistent with the contention that the pandemic lineage showed more reduced genetic diversity than the endemic lineage of the rice blast fungus (*60*). We speculate that the wheat blast fungus is still evolving rapidly, following the same evolutionary process as the loss of female fertility during the spread of the rice blast fungus.

Further analysis to identify and characterize additional genes responsible for female sterility, their functions, interactions, and phenotypic effects are required to better understand why and how *P. oryzae* lost female fertility during its evolution and the advantage conferred by the phenotype in nature. However, the present study opens new perspectives on the biological importance of loss of female fertility in plant pathogenic fungi. Our findings provide valuable information for management of plant disease before a pandemic.

## Materials and Methods

### Strains and culture conditions

*Pyricularia oryzae* isolates and transformants used and generated in this study were stored in our laboratory, with the mycelial plugs submerged in YEG-glycerol (5 g/L yeast extract, 20 g/L glucose, and 20% glycerol) at − 80°C. Originally, the female-fertile isolates CH598 and CH524 were collected from Yunnan, China by Dr. Li Chengyun, the female-sterile isolate Kyu89-246 from Japan by Dr. Shinzo Koizumi, and P2 (P-2) and Hoku-1 (Hoku1) from Japan by Dr. Tohru Teraoka (25). Strains were grown on oatmeal agar plates (25 g/L oatmeal flour, 2.5 g/L sucrose, and 1.5% agar) at 25°C for general maintenance. Crosses were performed by confrontationally placing two strains to mate on rice flour medium (20 g/L rice flour, 2 g/L yeast extract, and 1.5%agar) and grown for 4 weeks at 20°C under fluorescent light. At least five technical replicates were conducted for each cross.

### Progeny acquisition

After 4 weeks cultivation, mature perithecia were collected and crushed on glass microscope slides to observe ascospores. Ascospores were inoculated onto 1.5% agar plates and incubated overnight at 28°C. Single germinated ascospores were picked up with a fine glass needle and tweezer under a microscope and inoculated onto oatmeal agar plates. Only an ascospore per perithecium was collected to ensure the randomized acquisition of progenies.Mating type and knocked-in markers were determined by PCR (see Table S4) for every progeny obtained.

### Isolation of genomic DNA and HiSeq analysis

Mycelial plugs were individually inoculated into 5 mL YEG broth (5 g/L yeast extract and 20 g/L glucose) in Petri dishes and incubated for 5 days at 28°C without shaking. The culture medium was removed from the mycelia using paper towels, and the mycelia were frozen with liquid nitrogen in 1.5-mL centrifuge tubes. The frozen mycelia were ground in a pestle, and the powder was suspended in DNA Lysis Buffer [400 mM Tris-HCl (pH 7.5), 60 mM EDTA, 150 mM NaCl, and 350 mM SDS]. After centrifugation, the supernatant was purified with chloroform, and the DNA was precipitated with 0.3 M sodium acetate and 50% isopropanol.Pellets were dissolved in 50 µL of 20 µg/mL DNase-free RNase A solution and incubated at 65°C for 30 min. For HiSeq analysis, four monocultural DNA samples (25µL of 1 µg/µL) were mixed by phenotype, and then libraries were prepared using the TruSeq DNA PCR Free High Throughput Library Prep Kit (Illumina). Paired-end reads of 100 bp were generated with an Illumina platform (library preparation and sequencing were performed by Macrogen Japan).

Sequence reads were analyzed using QIAGEN CLC Genomics Workbench. First, raw reads were filtered using “Trim Reads” tool to discard nucleotides with error probabilities above 0.05, or with more than one ambiguous base. Then, the trimmed reads were mapped to the *P. oryzae* 70-15 version 8 (MG8) reference genome using the “Map Reads to Reference” tool.Reads of each phenotype were individually proceeded by following parameters: match score, mismatch cost, insertion cost, deletion cost, length fraction and similarity fraction were set at default (1, 2, 3, 3, 0.5 and 0.8, respectively). Paired distances were auto-detected, alignment was performed locally, and non-specific matches were discarded. Then variants between reads and the reference were detected by “Basic Variant Detection” after discarding reads non-specific, of broken pairs or with coverage > 1,000. These variants were then compared between the two phenotypes using “Compare Sample Variant Tracks” to retain inconsistent nucleotides as substitutions (detection frequency > 50%). Loci of the substitutions were visualized with SnapGene software (Insightful Science, available at https://www.snapgene.com) to explore the inherited region, *i*.*e*., FS1, in this research. In addition, contigs were assembled *de novo* for each phenotype using CLC Genomics Workbench to precisely determine the genotyping markers and candidate genes. Parameters were set as follows. Word size and bubble size were automatically determined at 23 and 50, respectively, paired distances were auto-detected, and scaffolds were created. Reads were then mapped back to contigs, where default costs and fractions were set as above. Contigs were updated by locally aligned reads.

### Isolation of total RNA and RNA-Seq analysis

Total RNA isolation was performed using the Monarch^®^ Total RNA Miniprep Kit (New England Biolabs). Mycelia were cultured in RY broth (20 g/L rice flour and 2 g/L yeast extract; boiled in a microwave oven, and the supernatant was recovered by centrifugation). For the starter culture, mycelia were grown on RY agar (RY broth supplemented with 1.5% agar) for 5 days at 28°C. Three mycelial plugs per sample were cut from colony edges with a cork borer (5 mm in diameter), placed in a triangle in 2 mL RY broth in Petri dishes (40 mm in diameter) and incubated without shaking for 7 days at 20°C under fluorescent light. Mycelia, treated as the DNA extraction procedure, were suspended in 1 × DNA/RNA Protection Reagent (New England Biolabs), and RNA Lysis Buffer (New England Biolabs) was added to the supernatant after centrifugation. Steps hereafter were conducted in accordance with the supplier’s protocol. The RNA was eluted in 100 µL nuclease-free water. For transcriptomic analysis, cDNA libraries were prepared with the TruSeq Stranded mRNA LT Sample Prep Kit (Illumina) and sequenced on an Illumina platform to generate paired-end reads of 150 bp (performed by Macrogen Japan).

Reads were analyzed using CLC Genomics Workbench. Briefly, raw reads were filtered using “Trim Reads”. Then, the reads were mapped to the MG8 reference genome using “RNA-Seq Analysis” tool, and the “Convert to Tracks” algorithm was used to extract gene track and calculate gene expression levels (RPKM values). Three progenies for each phenotype were treated as biological replicates. Principal component analysis and heat map creation were performed using the “PCA for RNA-Seq” and “Create Heat Map for RNA-Seq” tools, respectively. A heat map with complete cluster linkage was generated based on Euclidean distance without filtering (taking all transcripts into account).

### Transformation of *P. oryzae*

CRISPR/Cas9-mediated gene disruption, complementation, introduction of point mutations, and marker knock-in were performed as described previously (*38, 39*). pMK-bar was constructed by replacing the *HPH* cassette in pMK-dGFP (*38, 39*) with the *BAR* cassette. pMK-dGFP and pMK-bar were used as the backbone vectors. Pairs of flanking regions or a single genomic region with point mutations were cloned by PCR (the primers are listed in table S4), digested by appropriate restriction enzymes, and inserted into pMK-dGFP or pMK-bar. Target sites of CRISPR/Cas9 were designed and inserted as performed in previous studies (*38, 39*).

Transformants were verified by PCR and Southern blot analysis. At least double mutants were obtained from independent transformation procedures and tested for consistence of the phenotype.

### Asexual phenotypic analyses

Vegetative growth and asexual sporulation were performed on rice flour medium and CM (*61*). Mycelia were inoculated from the edges of 6-day-old starter cultures with bamboo sticks.After 6 days of incubation at 28°C, vegetative growth was measured. For conidial induction, aerial hyphae of 7-day-old cultures were brushed away, and plates were placed under a black light (FL15BLB, Toshiba) for 3 days at 25°C. Conidia were collected from the plates by pouring 5 mL distilled water (D.W.) and scrubbing the surface. The collected spores were precipitated by centrifugation (10,000 ×*g*, 5 min), resuspended in 500 µL D.W. and counted by a Thoma hemocytometer. Appressorium formation was induced by placing 20 µL of conidial suspensions (5 × 10^4^/mL) onto borosilicate 18 mm × 18 mm glass coverslips (Matsunami) followed by incubation under fluorescent light at 25°C for 6 hours. Invasive capacities were tested on onion scales boiled in a microwave oven (500 W and 30 sec). Rates of appressoria evading into onion cells were calculated 24 hours post-inoculation of the conidial suspensions. Conidial detachment assay was measured as previously (*12*), with slight modifications. Conidia-induced plates were stamped on 1.5% agar plates, and the conidia were recovered with 2 mL of D.W. Centrifuged conidia were resuspended in 20 µL D.W. and counted. All experiments except conidial detachment were performed in triplicate. The conidial detachment assay was repeated ten times.

### Analyzing deposited genome assemblies

Genome data were downloaded from NCBI Genome (https://www.ncbi.nlm.nih.gov/data-hub/genome/?taxon=318829), as well as DDBJ Sequenced Read Archive (https://ddbj.nig.ac.jp/search) under the accession numbers DRR059884–DRR059895. Fastq reads downloaded from DDBJ were assembled as HiSeq analysis. Local BLAST searches for *Pro1* ORF were carried out by CLC Genomics Workbench using the BLAST database generated from all the assemblies. Hits were exported in fasta and re-aligned to the query by SnapGene.Aligned sequences were individually assessed to find substitutions.

### Protein structure prediction

The 3D structures of Pro1 variants were predicted using ColabFold (*49*) to determine whether each Pro1 variant retained its function. The program is publicly available on Google Colab (https://colab.research.google.com/github/sokrypton/ColabFold/blob/main/AlphaFold2.ipynb), and is based on MMseqs2 (*62*). The monomer structures were predicted based on information from UniProt Reference Clusters and environmental databases, without PDB information as protein templates. The Amber programme (*63*) was not used. For each variant, the predicted rank_1 structure was compared with that of Pro1^CH598^ (variant #0) using PyMOL (https://pymol.org/2/). For variants whose Zn(II)_2_Cys_6_ DNA-binding domain was more than partially conserved, the domain was aligned with that of the variant #0 (cycles: 5 and cutoff: 2.0). For the others, the alignment was performed globally so that the maximum likelihood was achieved between all regions of the protein.

### Construction of phylogenomic and phylogenetic trees

Phylogenomic trees were constructed using IQ-TREE (*64*) with 1,000 replicates of ultrafast bootstrap (*65*). For the phylogenomic tree, genomic assemblies with Pro1 variants were analyzed using BUSCO (*66*) to detect and export BUSCO genes. Single-copy BUSCO genes commonly detected in all assemblies were collected, and multiple alignments were performed using Clustal Omega (*67*). These alignments were merged with the isolates, and a supermatrix was used for tree construction. For the Pro1 phylogenetic tree, multiple alignments were performed using the nucleotide sequence of *Pro1* variants (without an intron).

## Acknowledgments

We acknowledge Professor Elisabeth Fournier and Professor Didier Tharreau, Montpellier University, for helpful discussions and advice. We thank Robert McKenzie, PhD, from Edanz (https://jp.edanz.com/ac), for editing a draft of this manuscript.

## Data, Materials, and Software Availability

*Pyricularia oryzae* gene data are available in GenBank under the following accession numbers: *Pro1*, MGG_00494; *Mak1*, MGG_04943; *Mak2*, MGG_09565; *Mek1*, MGG_06482; *Pro40*, MGG_01636; *Nox1*, MGG_00750; *Nox2*, MGG_06559; *Pro41*, MGG_09956; *Pls1*, MGG_12594; *Ham-5*, MGG_06673; *Ham-7*, MGG_10588; *Ham-8*, MGG_01593; *Ham-9*, MGG_00492; *EsdC*, MGG_12316.

## Author contributions

Designed research: TARA, TKA; Performed research: MU, TKO, AF, KK, TARA; Contributed new reagents/analytic tools: KK, TARI, TT; Analyzed data: MU, TKO, AF, TARI, TT, TARA, TKA; Wrote the paper: MU, TARA, TKA.

