## Supplementary Materials for "Why do some fungi want to be sterile? The role of dysfunctional Pro1 in the rice blast fungus"

**This PDF file includes:**

Figs. S1 to S11

Tables S1 to S4

Fig. S1. Mating capability of six field isolates of *Pyricularia oryzae*. (A) Perithecia developed by crossing CH598×CH524, CH598×Kyu89-246, CH598×P2, and Hoku-1×CH524. Hoku-1, Kyu89-246, and P2 did not develop perithecia. (B) Asci and ascospores developed by crossing CH598×CH524 and CH598×Kyu89-246. Perithecia contained no mature asci or ascospores between CH598×P2 nor Hoku-1×CH524. Bars indicate 50 µm.

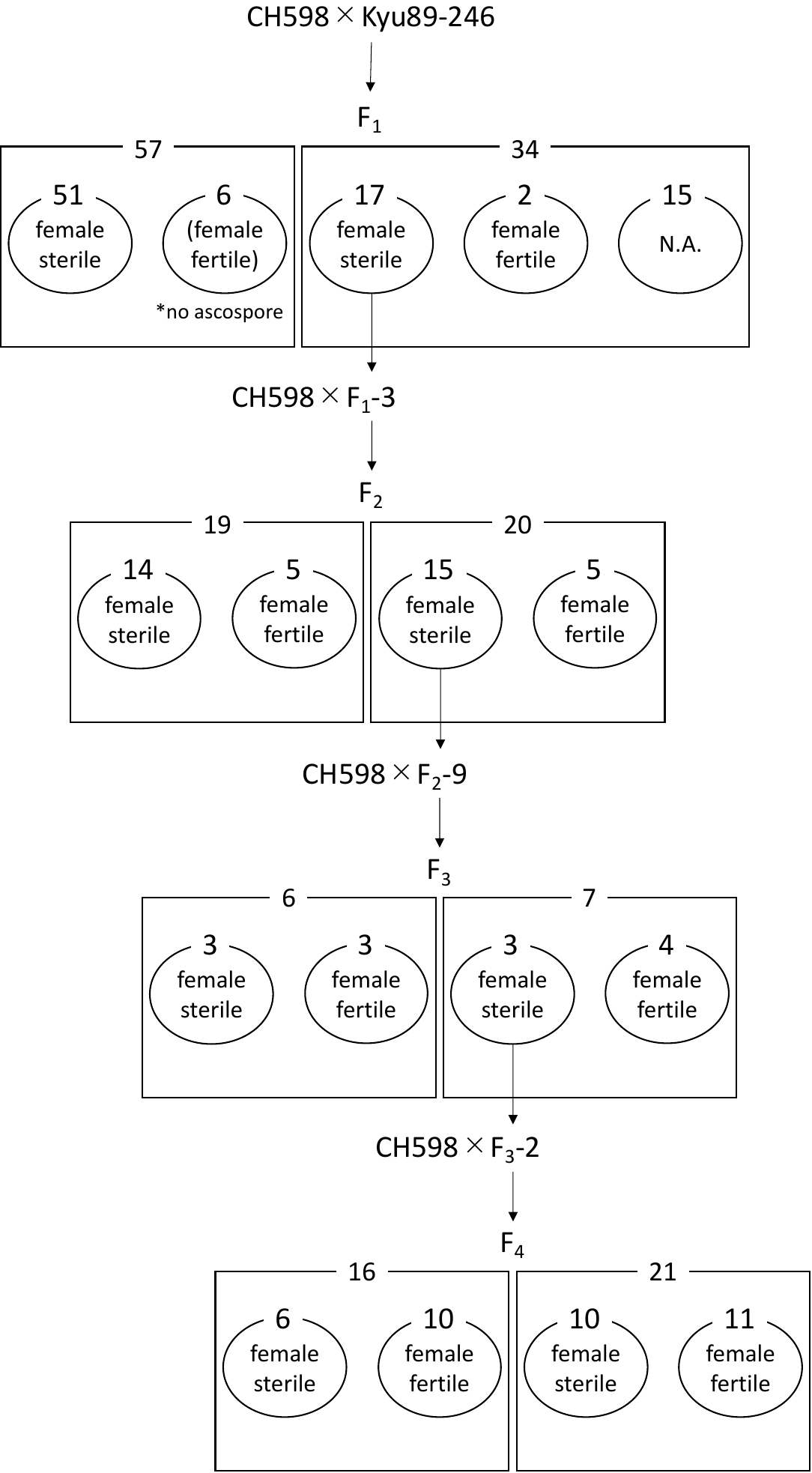

Fig. S2. Schematic diagram and the number of progenies obtained. Female-sterile progenies in each generation were crossed with CH598, and the mating-type and female fertility of the progenies were examined. N.A., not assessed.

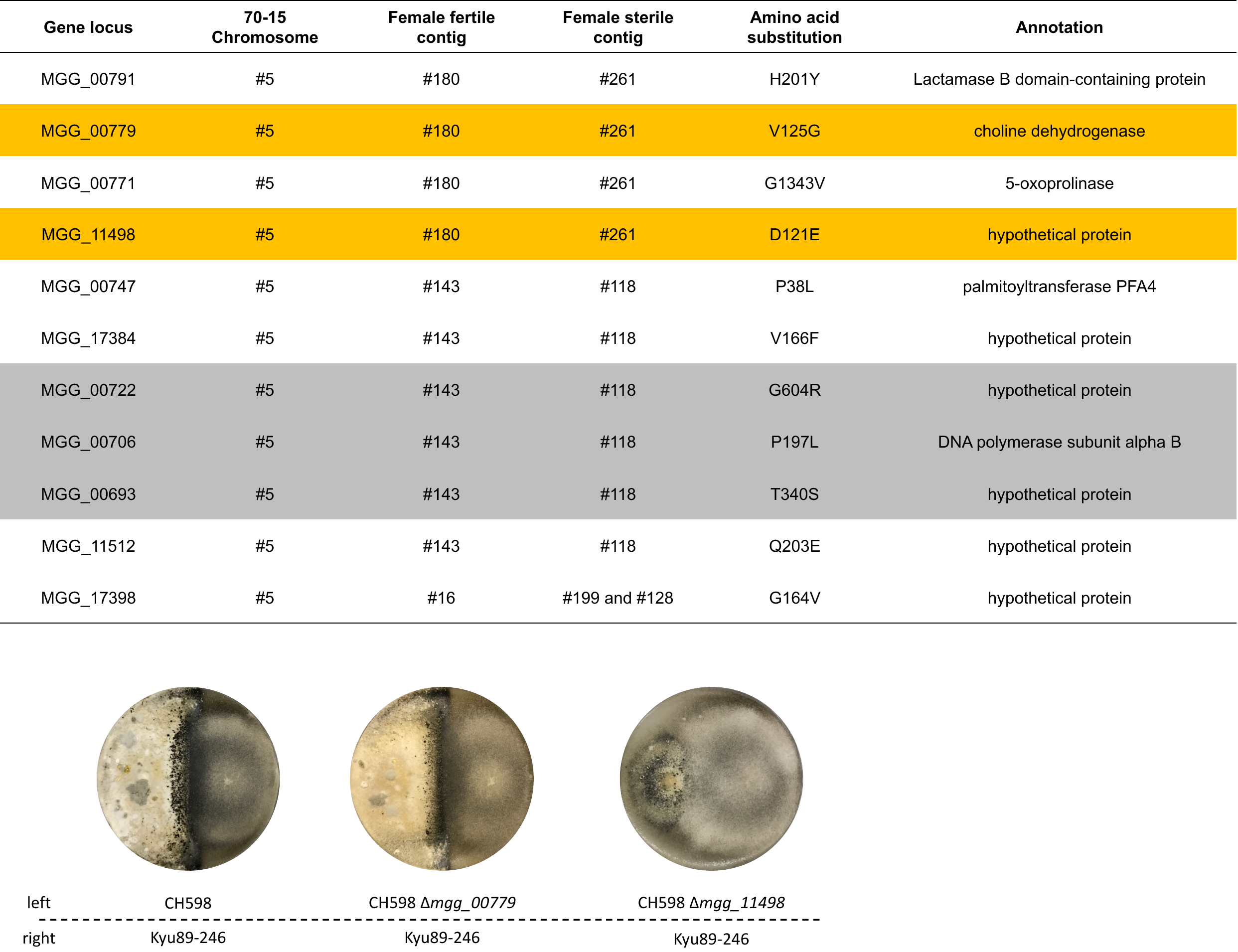

Fig. S3. Candidate genes involved in loss of female fertility in the FS1 region. The candidate genes containing amino acid substitutions were listed in the upper table. The two genes were related to perithecium formation (shaded in orange in the upper table and lower pictures) and the three genes (colored with grey) were predicted to be lethal genes.

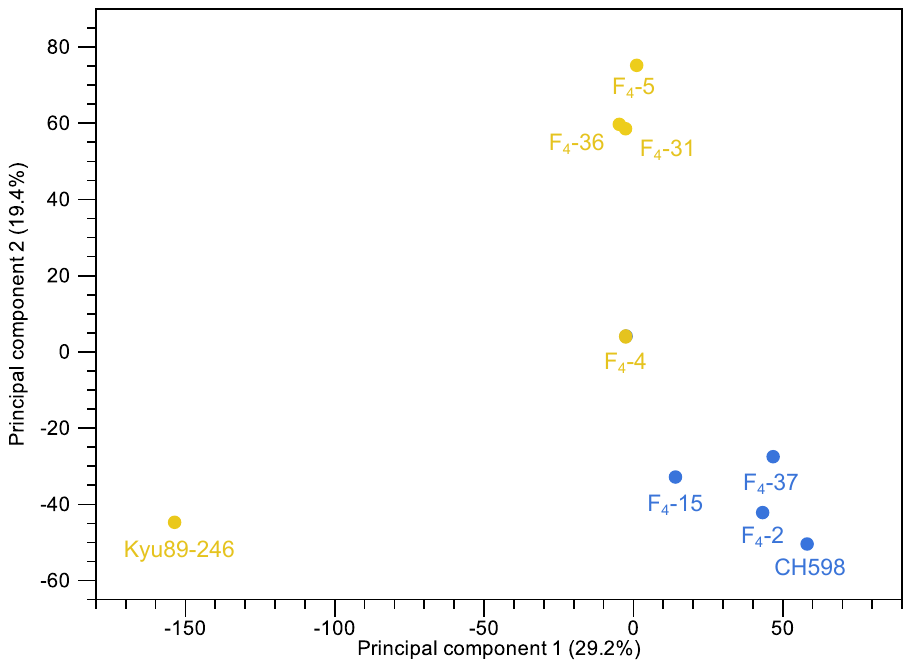

Fig. S4. Principal component analysis of RNA-seq data including F_4_-4. Female-sterile-evolved F_4_-4 showed relatively closer expression patterns to female-fertile strains.

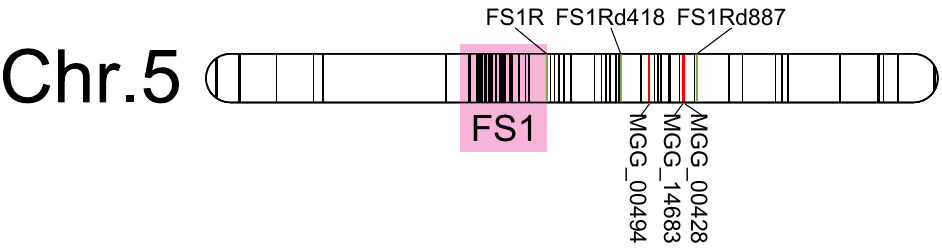

Fig. S5. Refined mapping of nucleotide substitutions. The reads of female-fertile progenies were re-aligned to the *de* *novo* assembled genome of female-sterile progenies, and the detected substitutions were mapped to the reference genome.

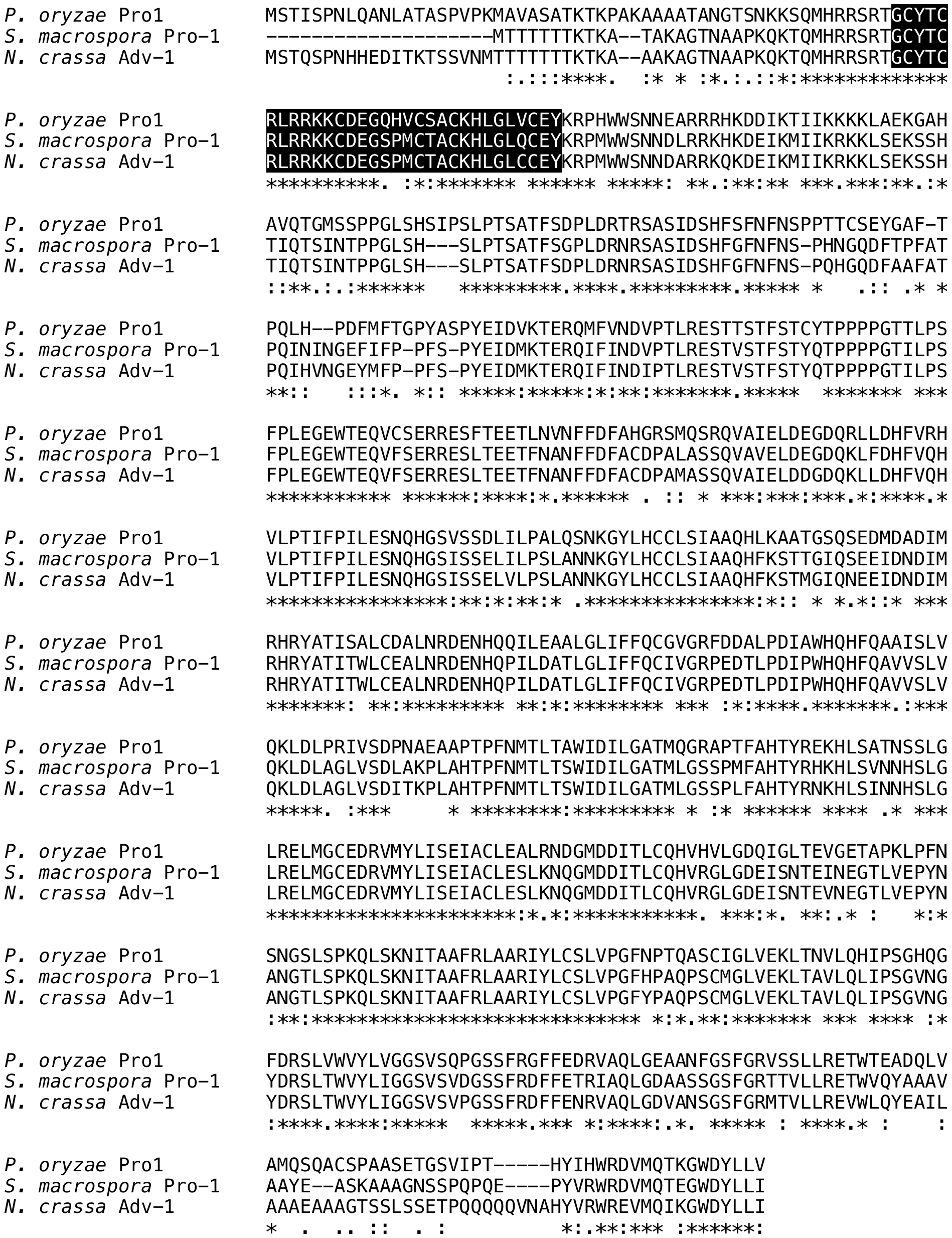

Fig. S6. Alignment of amino acid sequences of *P.* *oryzae* Pro1 and its orthologs in *Sordaria* *macrospora* and *Neurospora* *crassa*. The Zn(II)_2_Cys_6_ DNA-binding motif is shaded in black. Asterisks, double dots, and single dots represent consistent, similar, and dissimilar amino acids among the three species, respectively.

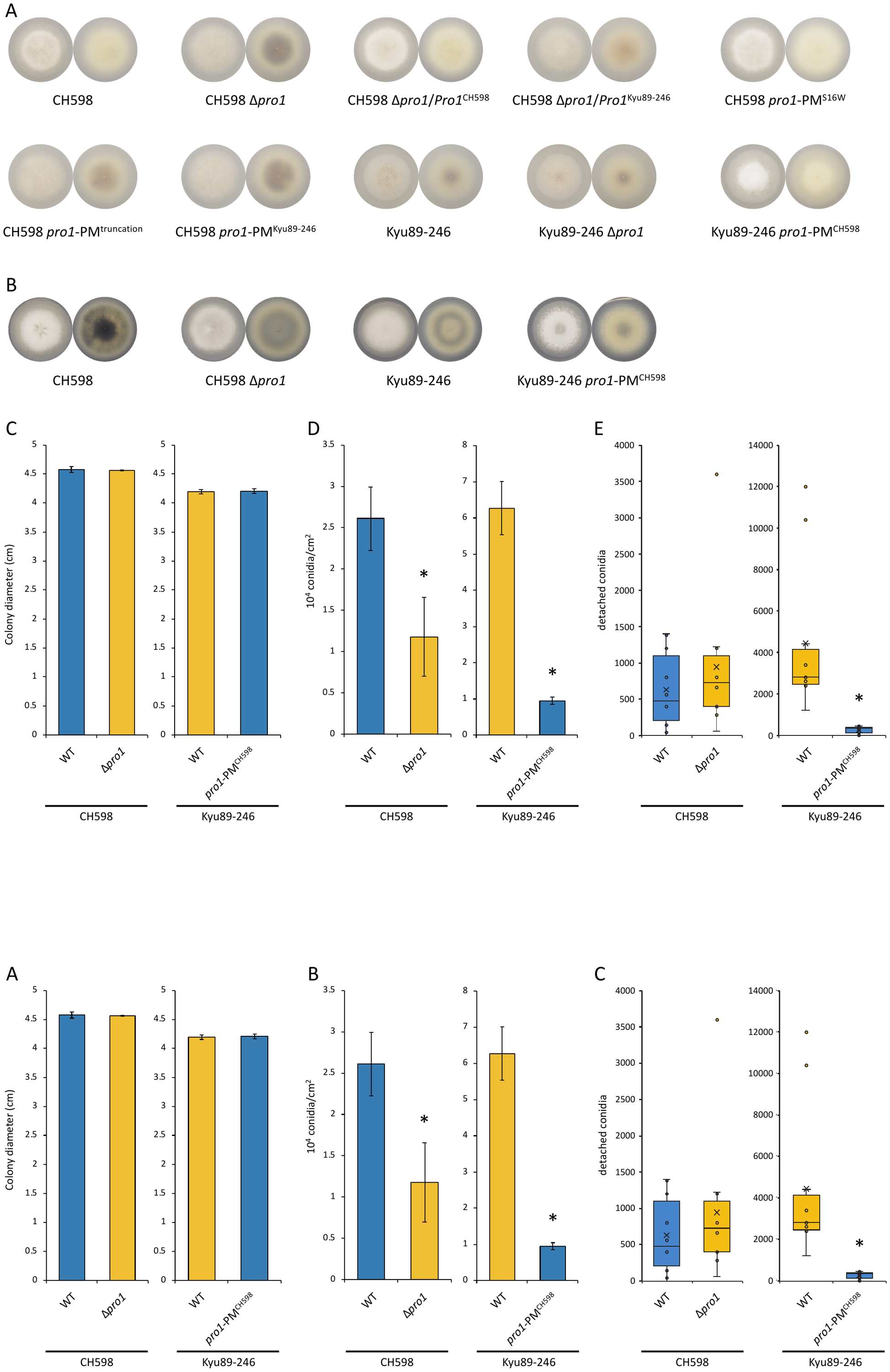

Fig. S7. Phenotypes associated with Pro1 function. Mycelial morphologies of CH598, Kyu89-246, and their transformants grown on (A) rice flour medium and (B) complete medium (CM) at 28°C for 6 days. Left, from the top; right, from the bottom. (C) Colony diameter, (D) conidial production, and (E) conidial detachment on CM. Bars and error bars in (C) and (D) indicate the mean ± SEM. Crosses and lines in (E) indicate the mean and median, respectively. Blue bars and boxes indicate the results of strains possessing the functional Pro1. Yellow bars and boxes indicate the results of strains possessing the dysfunctional Pro1. * *p* < 0.05 (Welch’s *t*-test).

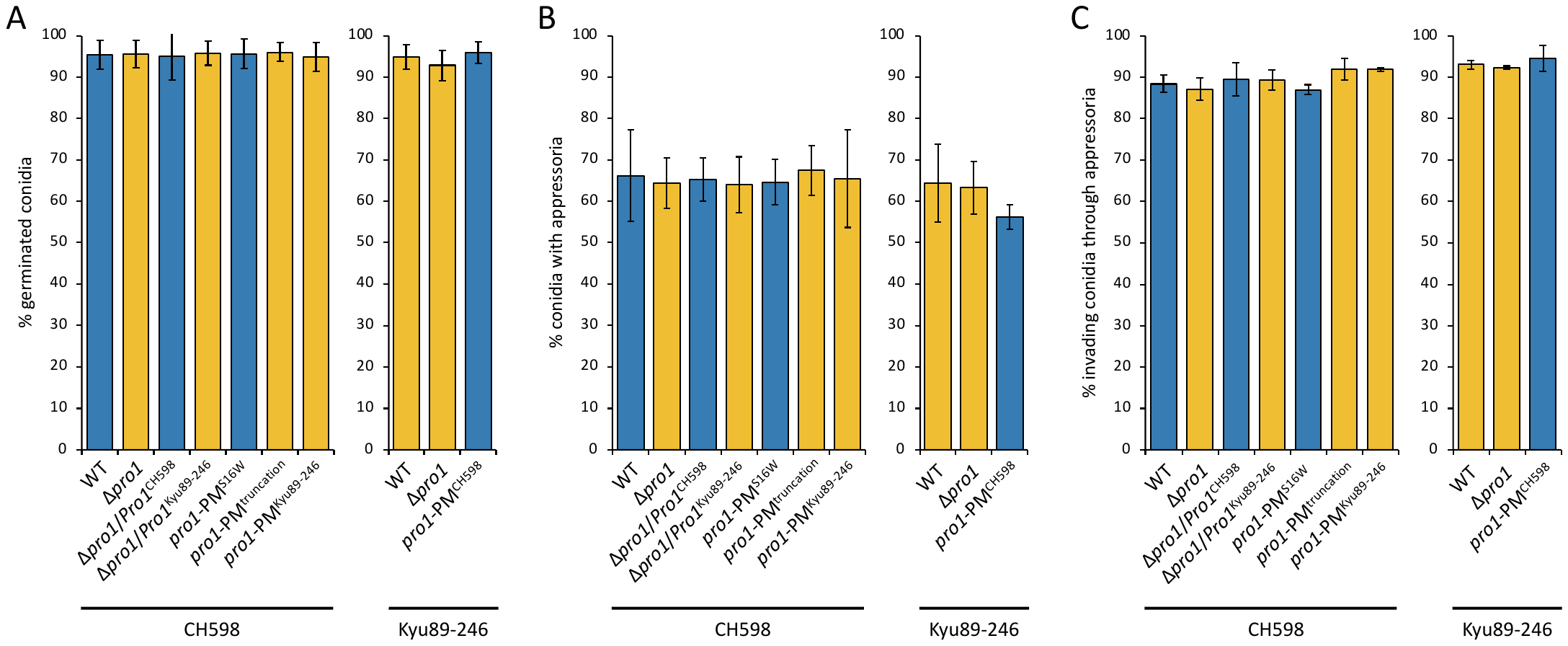

Fig. S8. Asexual phenotype associated with Pro1 function. (A) Percentage conidial germination and (B) appressorium formation, and (C) invasion through appressoria in the CH598 and Kyu89-246 genetic backgrounds. No significant difference was detected in each genotype. Bars and error bars indicate the mean ± SEM. Blue, strains with functional Pro1; yellow, with dysfunctional Pro1.

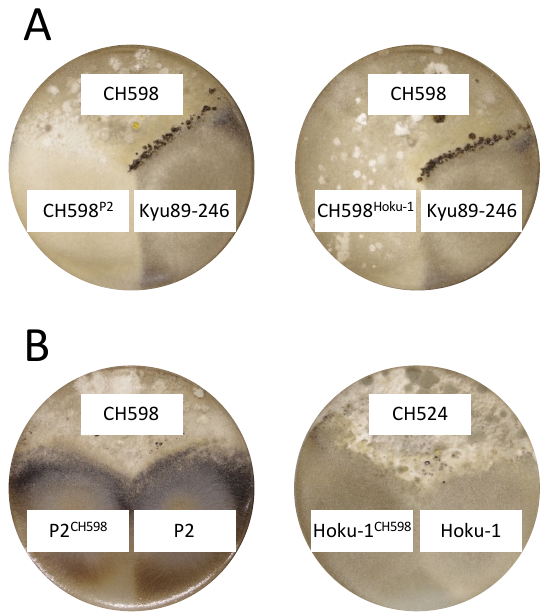

Fig. S9. Functions of mutated Pro1 in P2 and Hoku-1 as determined by crossing. CH598 mutants possessing P2- or Hoku-1-derived *Pro1* variants did not develop perithecia. Functional *Pro1* integrated into P2 and Hoku-1 did not rescue their fertility. Superscripts represent the sequence origin of *Pro1*.

Fig. S10. Predicted protein structures of Pro1 variants. The rank_1 predictions of protein folding by ColabFold are shown. The domain colored yellow in variant #0 is the Zn(II)_2_Cys_6_ DNA-binding domain. Protein structures for each variant are colored red, together with variant #0, colored light grey. Structures were aligned at the DNA-binding domain, if possible, or oriented most plausibly.

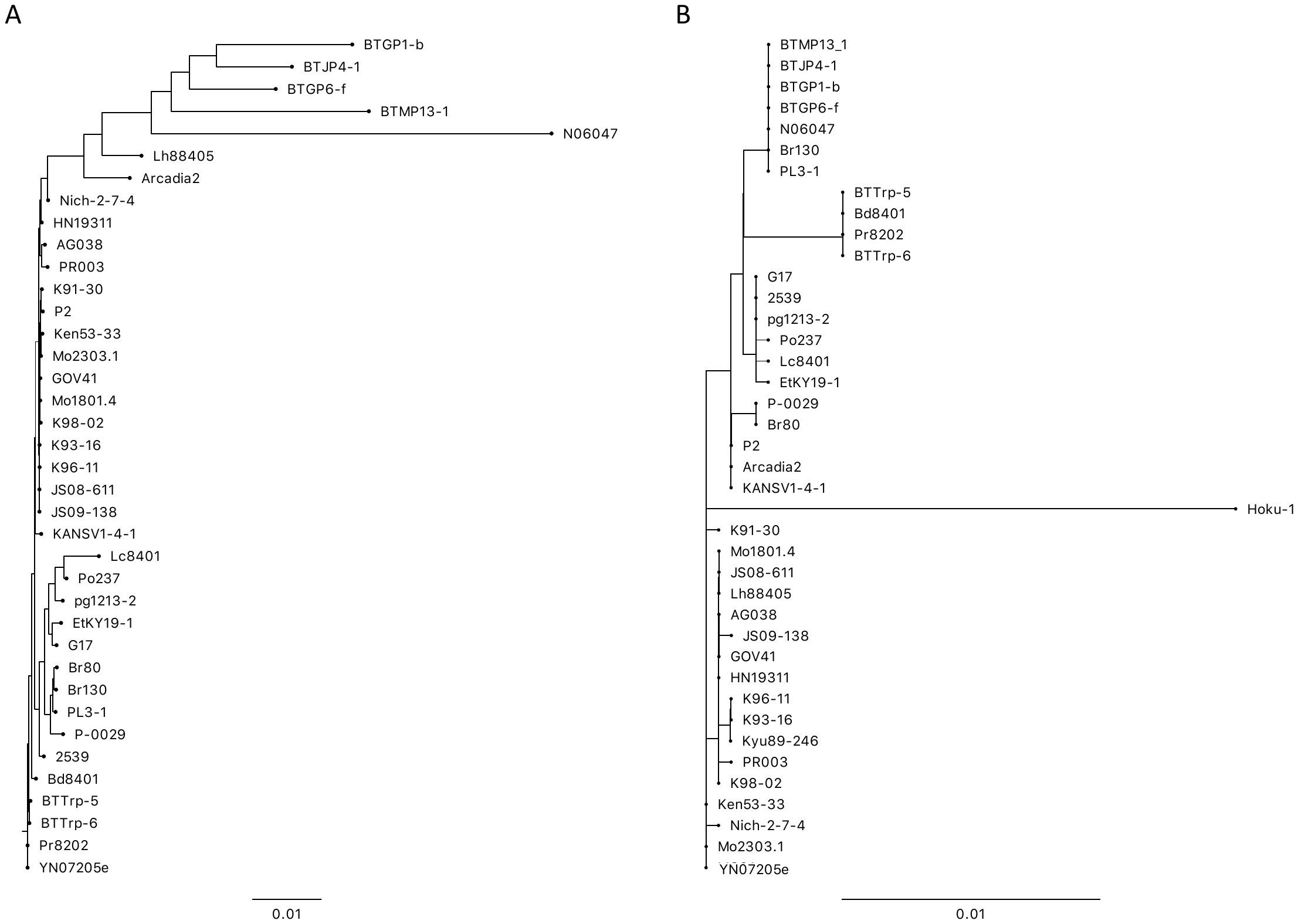

Fig. S11. Phylogenomic and *Pro1* phylogenetic trees. (A) A phylogenomic tree of 38 isolates drawn by nucleotide sequences of 468 BUSCO genes, which were detected in all strains as single copies. (B) A phylogenetic tree of *Pro1* ORF sequences from the 40 isolates possessing Pro1 mutations. Kyu89-246 and Hoku-1 are missing in the phylogenomic tree because of no available genomic sequence. YN07205e, a Yunnan (the origin of *P.* *oryzae*) isolate possessing functional Pro1.

**Table S1. Segregation of mating type and female fertility in F_4_ progenies.** The numbers of strains for each genotype or phenotype are shown. Numbers in Parentheses are the values when F_4_-4, which is thought to be a putative female-sterile-evolved strain because of inconsistent genotype through FS1 region, is treated as a female-fertile strain.

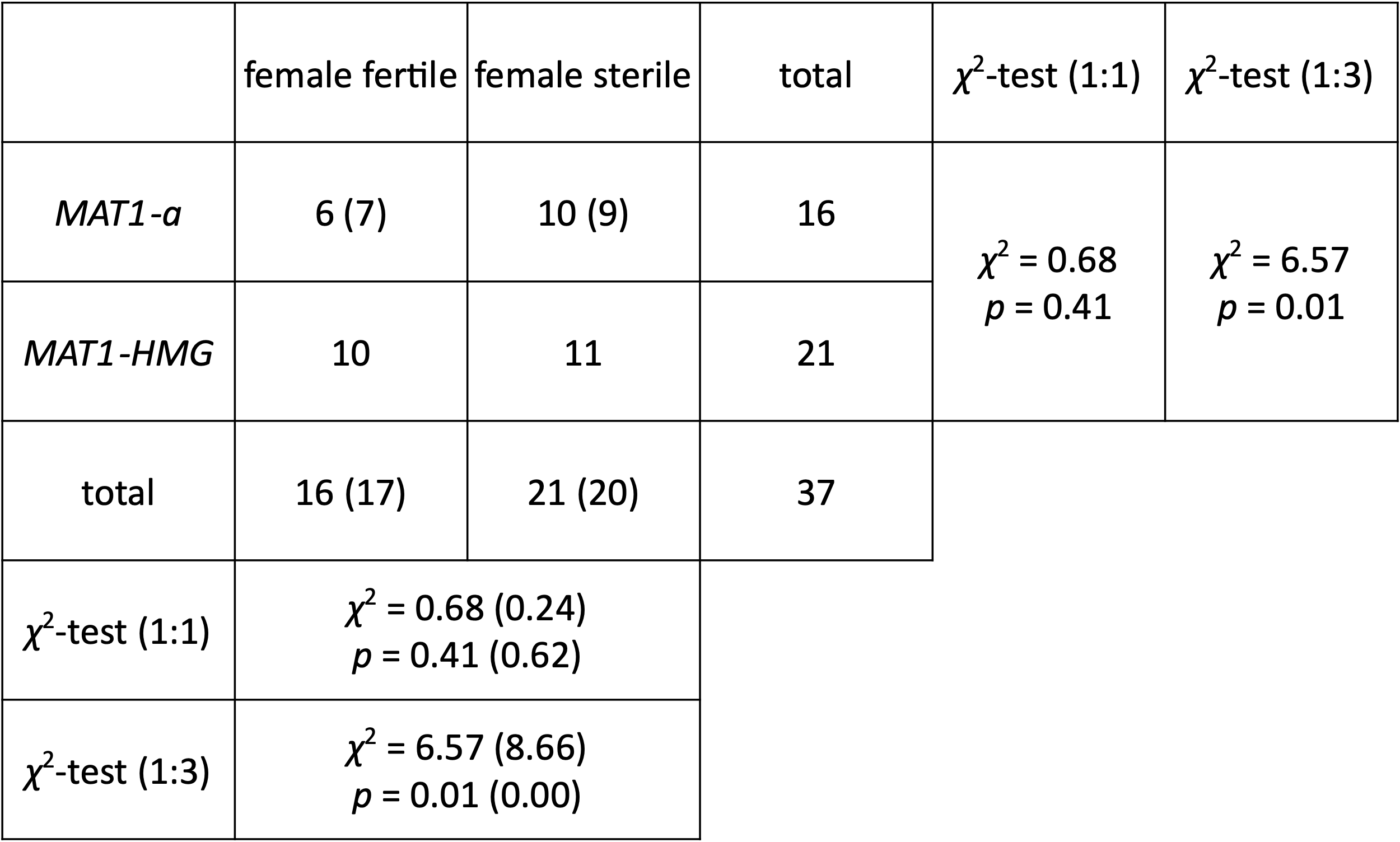

Table S2. Genes upregulated or downregulated by Pro1.

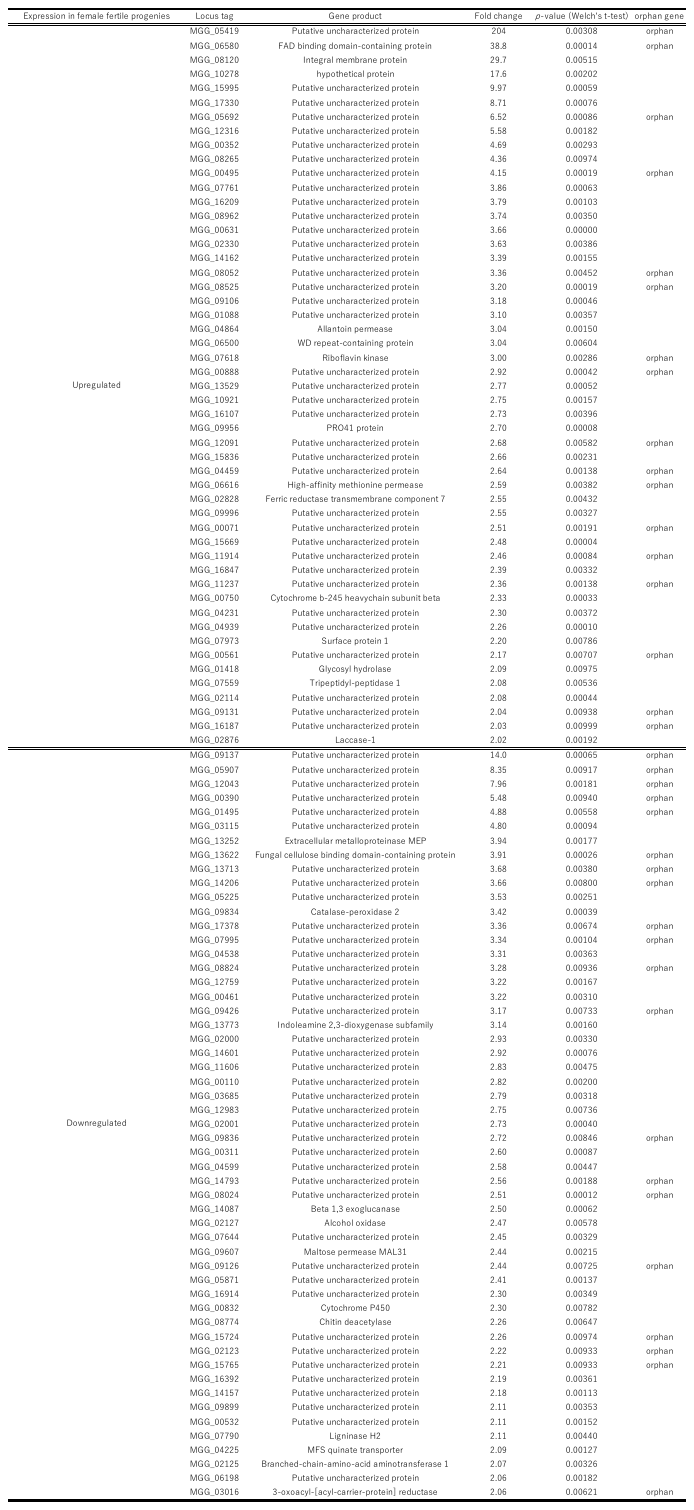

Table S3. Isolates and detected Pro1 mutations. BLAST search against the *Pro1* open reading frame (ORF) was carried out, resulting in (A) assemblies containing the complete sequence of *Pro1* without any mutations, (B) assemblies including the entire *Pro1* sequence with more than one mutation, and (C) assemblies lacking the total ORF of *Pro1*. Strains colored red is redundant in the table.

A

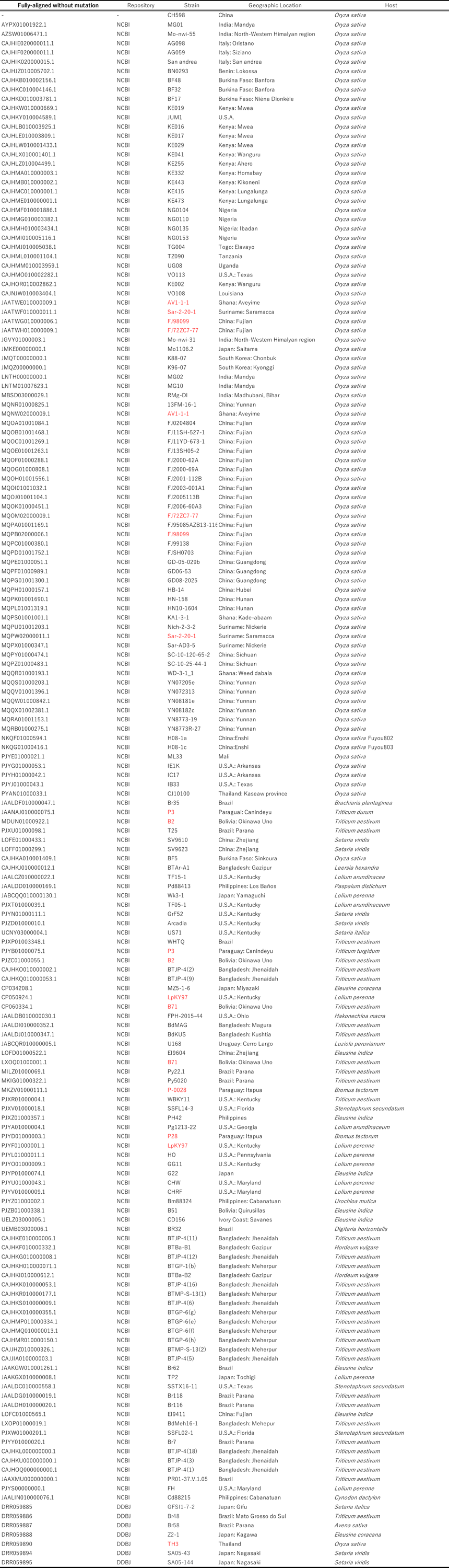

(Continued from the previous page)

B

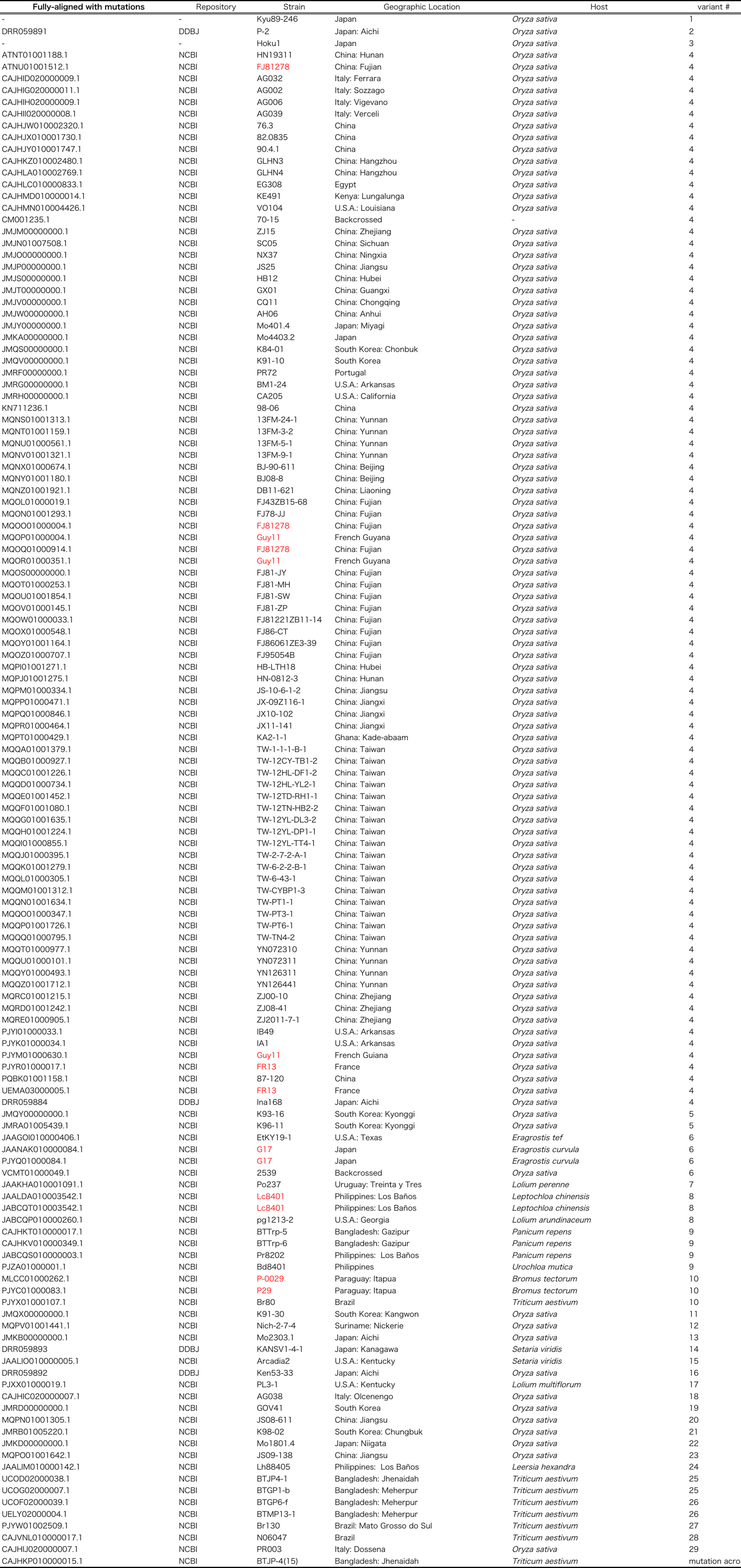

C

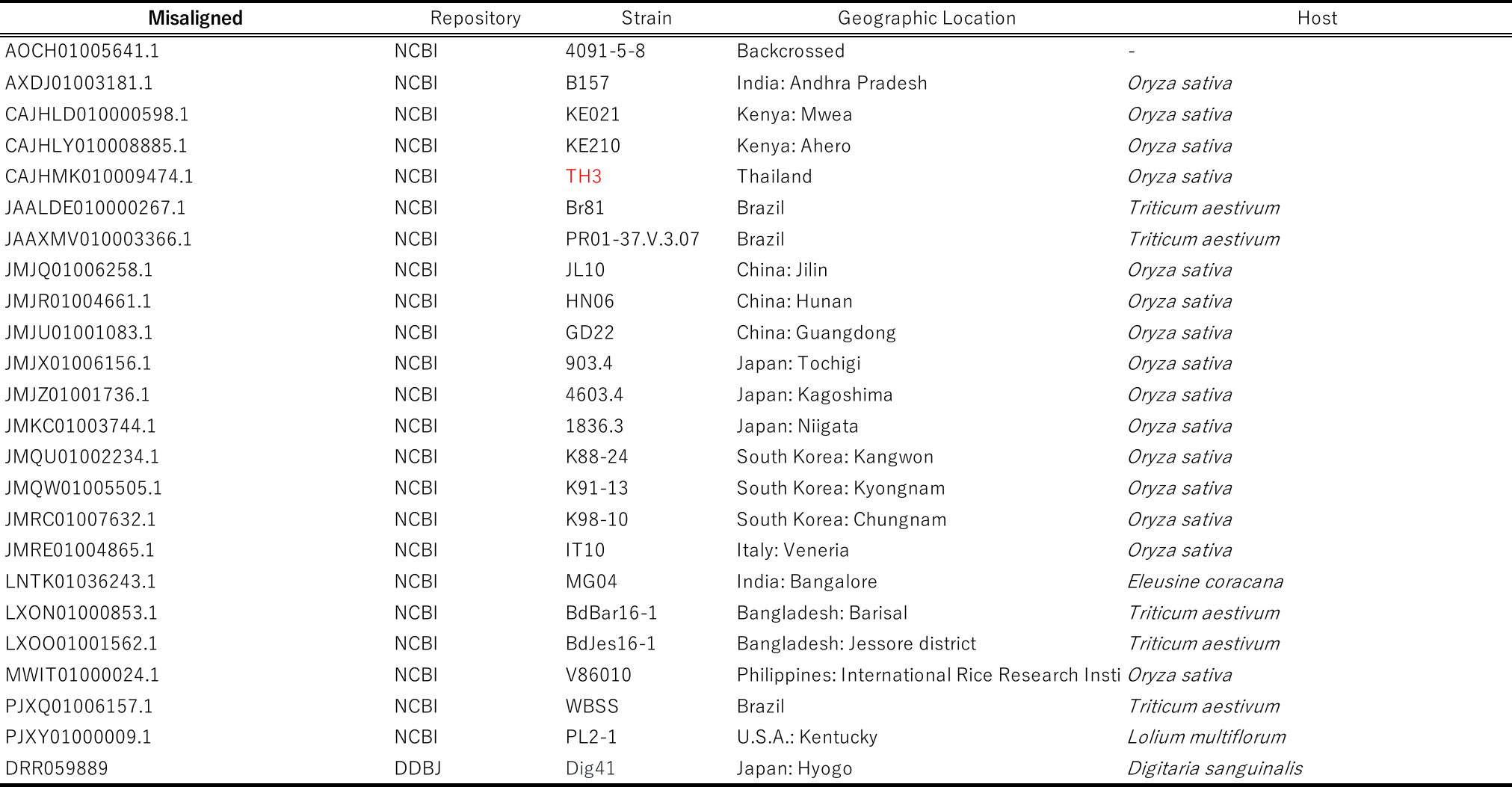

Table S4. Primers used in this study.

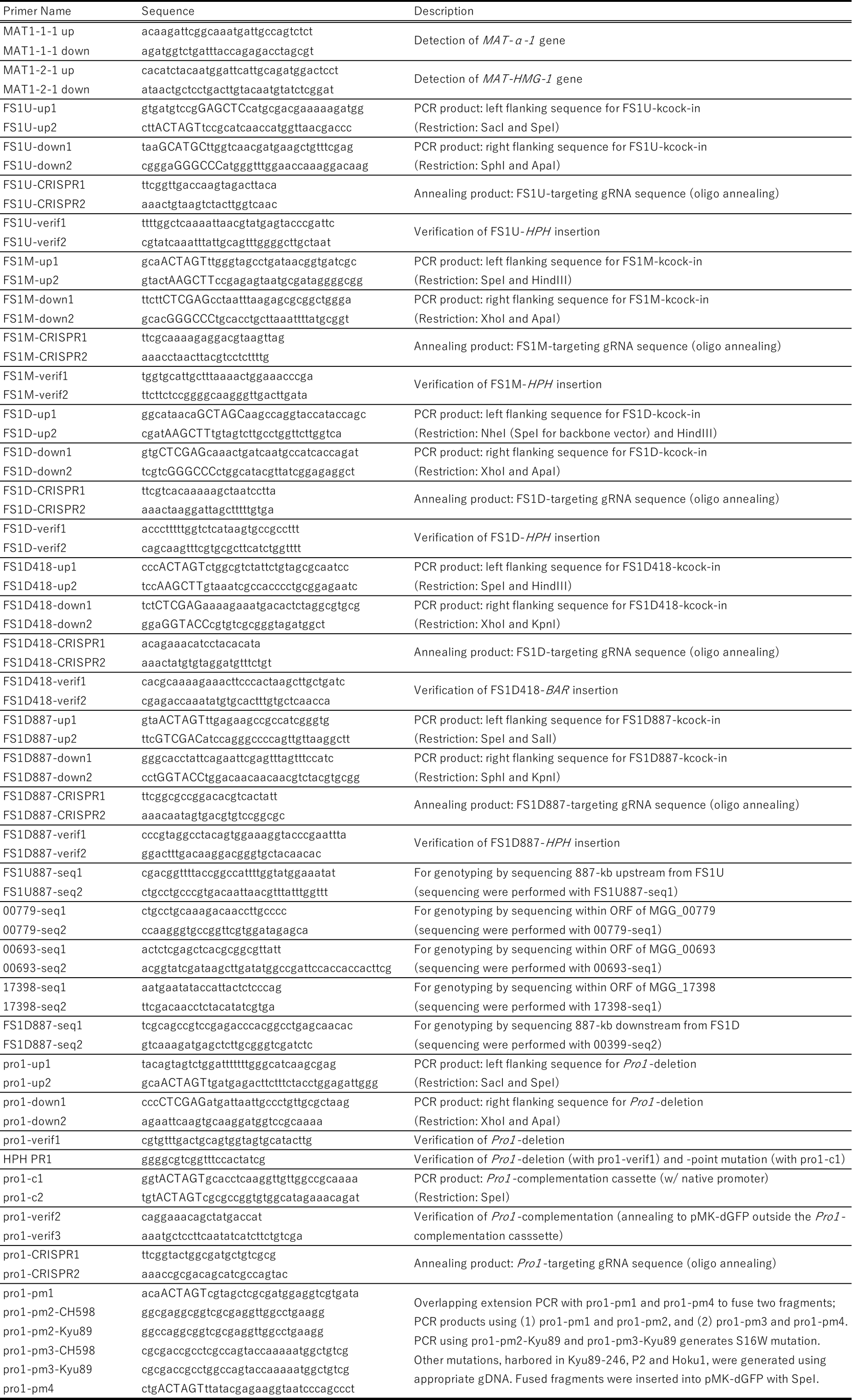
